# Direct cell-to-cell transport of Hedgehog morphogen is aided by the diffusible carrier Shifted/DmWif1

**DOI:** 10.1101/2025.06.04.657875

**Authors:** Carlos Jiménez-Jiménez, Gustavo Aguilar, Clara Fernández-Pardo, Markus Affolter, Isabel Guerrero

**Author notes:** Equal contribution (Carlos Jiménez-Jiménez, Gustavo Aguilar).

## Abstract

The establishment of morphogen gradients is fundamental during embryonic patterning. Yet, the mechanisms by which morphogens disperse remain highly controversial. In a paradigmatic example, the dispersal of the morphogen Hedgehog (Hh) is both limited by and dependent on its lipid modifications. To address this paradox, several competing transport models have been proposed, ranging from lipid-shielding solubilizing factors to transport via specialized filopodia called cytonemes. In this study, we uncover that the lipid-shielding protein DmWif1/Shifted (Shf) participates in the contact-dependent cell transfer of Hh. Using nanobody-based immobilization, we demonstrate the tight association of Shf and Hh to cytoneme membranes. We show that Shf is anchored to membranes by the Hh co-receptor Interference Hedgehog (Ihog) and that the Ihog-Shf complex is required for Hh dispersal and signaling. Together, our results unify cytoneme-mediated transport and membrane release into a single dispersal model, in which local solubilization is required for cytonemes to exchange Hh-bound Shf at cell-cell contacts between presenting and receiving cytonemes.

## Introduction

Morphogen gradients are crucial for animal development, orchestrating the spatial organization of tissues (Rogers and Schier, 2011). Among them, the Hedgehog (Hh) family of proteins is essential for tissue patterning and the maintenance of tissue homeostasis after birth (Briscoe and Thérond, 2013). Disruption of the Hh signaling pathway is associated with pathologies such as holoprosencephaly, polydactyly, and several types of cancer (Zhang and Beachy, 2023).

The Hh protein is synthesized as a precursor of approximately 45 kDa and undergoes autocatalytic processing in the endoplasmic reticulum, resulting in two protein fragments (Hh-C and Hh-N) (Chen et al., 2011; Lee et al., 1994). The Hh-N fragment (hereafter referred to as Hh) undergoes two lipid modifications: a cholesterol molecule attached to its C-terminal end, and a palmitic acid added to a cysteine residue close to its N-terminus (Pepinsky et al., 1998; Porter et al., 1996). These lipid modifications strongly associate Hh with cell membranes, thereby limiting its ability to freely move through the extracellular environment (Peters et al., 2004). Counter-intuitively, these lipid moieties are also required for long-distance Hh signalling (Chamoun et al., 2001; Callejo et al., 2006; Gallet et al., 2006). In the past two decades, several models have been proposed to address this contradiction, such as the formation of multimers (Feng et al., 2004; Goetz et al., 2006), and transport via exovesicles (Gradilla et al., 2014; Matusek et al., 2014; Wang et al., 2022) or cytonemes (Bischoff et al., 2013; Hall et al., 2024, 2021; Sanders et al., 2013).

Cytonemes are actin-based cellular protrusions specialized in intercellular communication (González-Méndez et al., 2019; Ramírez-Weber and Kornberg, 1999; Zhang and Scholpp, 2019). In *Drosophila*, it has been demonstrated that Hh is located at cytonemes in various developing tissues (Bischoff et al., 2013; Callejo et al., 2011; Rojas-Ríos et al., 2012; García-Arias et al., 2025). The extension of these cellular protrusions correlates with the gradient of Hh signalling in the wing imaginal disc and in the abdominal histoblast nests, supporting their functional role (Bischoff et al., 2013; Aguirre-Tamaral and Guerrero, 2021;). Cytonemes can be produced by both signal-sending and signal-receiving cells, with the morphogen exchanged at contact sites between cytonemes, in a process reminiscent of neuronal synapses (Chen et al., 2017; González-Méndez et al., 2017; Huang et al., 2019; Roy et al., 2014). Yet, the mechanism by which Hh is exchanged at cell-cell contacts remains unknown.

We have previously identified Shf, a secreted protein that can disperse long-range and that interacts specifically with lipidated Hh (Glise et al., 2005; Gorfinkiel et al., 2005). Shf is orthologous to the vertebrate Wnt inhibitory factor (Wif1) protein (Hsieh et al., 1999), which regulates Wnt but not Sonic Hh dispersion (Sánchez-Hernández et al., 2012, Avasenov et al., 2012). The Wif1 family consists of a WIF-type box with a high affinity for lipids (Liepinsh et al., 2006; Malinauskas et al., 2011), and five EGF repeats (Glise et al., 2005; Gorfinkiel et al., 2005). Shf is critical for Hh gradient formation in *Drosophila*, and its absence restricts Hh signalling gradient to a single row of cells adjacent to the Hh source (Glise et al., 2005; Gorfinkiel et al., 2005). These results, together with the high diffusibility of Shf, led to the initial proposal that Shf would solubilize Hh, aiding the formation of the gradient through a diffusion mechanism.

Like Shf, Ihog and its homolog Brother of Ihog (Boi) are also critical for the maintenance of extracellular Hh levels (Bilioni et al., 2013; Callejo et al., 2011; Yan et al., 2010). Ihog and Boi have been described as Hh co-receptors (Zheng et al., 2010) but only Ihog is required for the long range Hh gradient (Simon et al., 2021). Consistent with this, overexpression of Ihog, but not Boi, regulates cytoneme dynamics and orientation (Aguirre-Tamaral and Guerrero, 2021; Simon et al., 2021; Yang et al., 2021; Aguirre-Tamaral et al., 2022).

In this study, we have revised the role of Shf during Hh transport and discovered that both Hh and Shf are tightly linked to cell membranes. Furthermore, we demonstrate that Shf is associated with membranes via the Ihog, which binds to Shf through its Ig domains. We also demonstrated that the Ihog-Shf complex is responsible for mediating Hh signalling. We propose that local solubilisation of lipid-modified Hh by Shf is required for cytonemes to exchange Hh at cell-cell contacts between presenting and receiving cytonemes.

## Results

### Retention patterns of immobilized artificial ligands

To distinguish between the different models of morphogen dispersal in polarized epithelia, we hypothesized that immobilizing a morphogen on the surface of recipient cells would reveal distinct retention patterns depending on the ligand’s effective membrane association. Specifically, whether it disperses anchored to producing membranes, as proposed by the cytoneme model, or freely diffuses through the tissue. To that end, we employed VHH4-CD8:mCherry (hereinafter “morphotrap”) (Harmansa et al., 2015), a membrane-tethered anti-GFP nanobody that when expressed in the wing disc epithelia, distributes evenly across the apical and basolateral plasma membranes and efficiently immobilizes GFP-tagged extracellular proteins on the cell surface (Harmansa et al., 2017, 2015; Matsuda et al., 2022).

We initially tested the immobilization assay using synthetic ligands with different modes of membrane association, which are expected to influence effective membrane tethering and 2D binding behavior (Hu et al., 2015). Using the *UAS*/Gal4 and LexO/LexA expression systems, we simultaneously expressed a secreted form of GFP in the posterior (P) compartment and morphotrap in the anterior (A) compartment (scheme in Figure 1A). Secreted GFP homogeneously decorated morphotrap-expressing cells throughout the apico-basal axis of the epithelium (Figure 1B-B’’’), with only a modest apical enrichment, indicating that fully soluble ligands disperse broadly in the wing disc and exhibit no clear localization bias. We then tested the retention pattern of a membrane-bound GFP (GFPext-CD8) in the same experimental configuration (scheme in Figure 1C). Apical and basolateral imaging revealed juxtacrine contacts between compartments (Figure 1D-D’’’). Most remarkably, a network of filopodial protrusions became apparent at the basal side of the epithelia (Figure 1D’’, E, E’, Supplementary Figure 1A). However, this setup did not result in the stabilization of any apical protrusions. The basal network extended across the AP compartment boundary, in some cases over tens of micrometers (Supplementary Figure 1A). Interestingly, the immobilized filopodia were not of regular thickness, presenting frequent thickenings along their length (Supplementary Figure 1A). We and others have previously reported basal filopodia in the wing disc (Bischoff et al., 2013; Callejo et al., 2011; Chen et al., 2017; González-Méndez et al., 2017; Yan et al., 2021); however, their visualization in fixed wing disc samples had so far only been possible upon expression of stabilizing proteins (Bischoff et al., 2013; Callejo et al., 2011).

**Figure 1.**
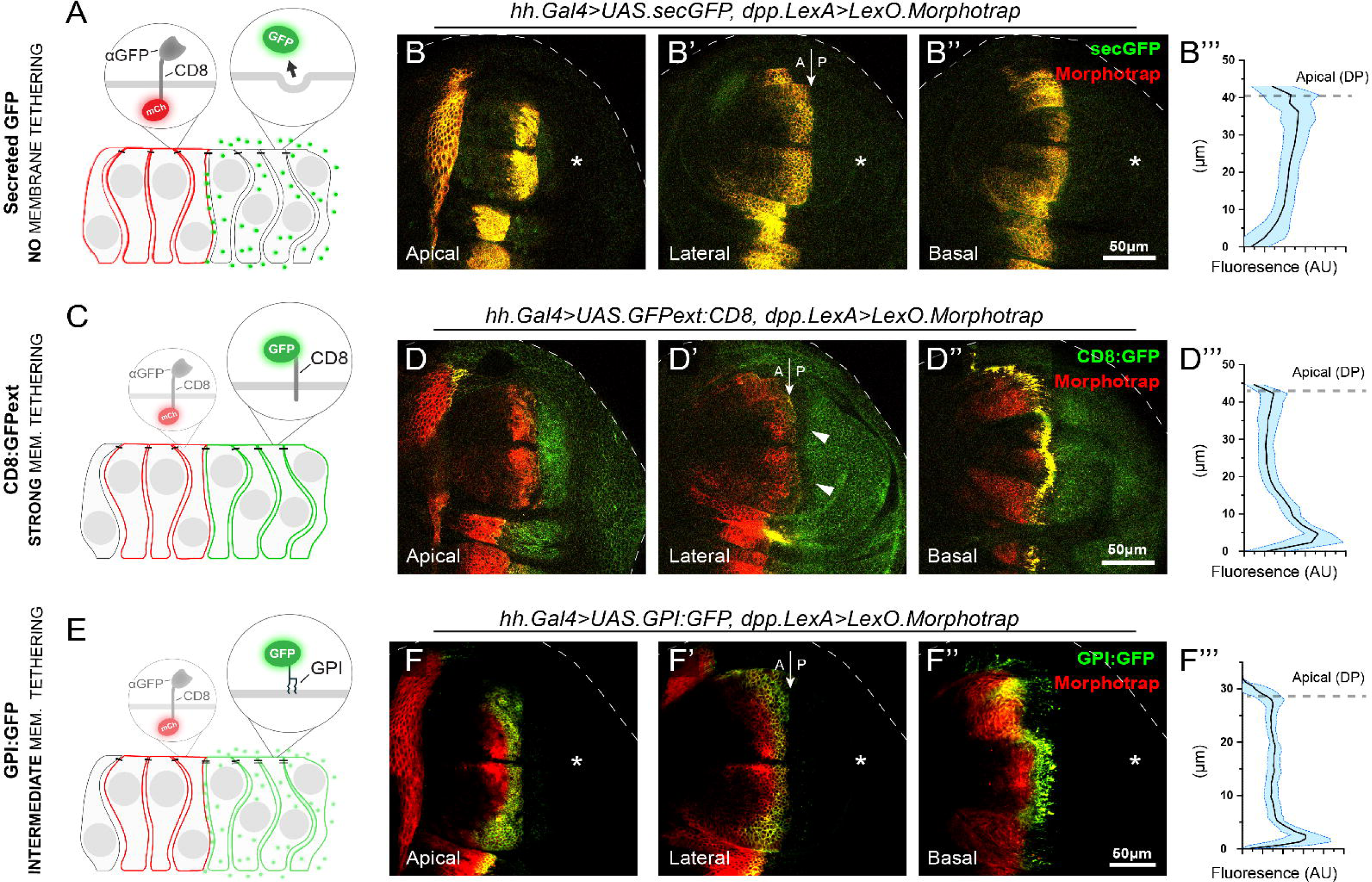
Nanobody-based strategy to immobilize artificial ligands. **A)** Scheme of morphotrap and secreted GFP expression domains in the wing disc. **B–B’’)** Wing disc expressing morphotrap in Hh-receiving cells (dpp.LexA) and secreted GFP in Hh-producing cells (hh.Gal4). Secreted GFP accumulates along the apico-basal axis of morphotrap-expressing cells. GFP signal in producing cells is much lower under these imaging conditions and is not visible (asterisk). **B’’’** shows apico-basal quantification of GFP fluorescence in receiving cells only (see Materials and methods). **C)** Scheme of morphotrap and membrane-bound GFP (GFPext-CD8) expression domains. **D–D’’)** Wing disc expressing morphotrap (dpp.LexA) and GFPext-CD8 (hh.Gal4). GFPext-CD8 accumulates at A/P contact sites, a prominent network of basal contacts is detected between A and P compartment cells. Both Morphotrap and GFPext-CD8 are reduced from the lateral membrane near the border (arrowheads). **D’’’** shows apico-basal quantification of GFP fluorescence in receiving cells. **E)** Scheme of morphotrap and GPI-anchored GFP (GPI:GFP) expression domains. **F–F’’)** Wing disc expressing morphotrap (dpp.LexA) and GPI:GFP (hh.Gal4). GPI:GFP accumulates at A/P contact sites and along the apico-basal axis of morphotrap-expressing cells. Signal in producing cells is much lower and not visible at these laser settings (asterisk). **F’’’** shows apico-basal quantification of GFP fluorescence in receiving cells.

We also tested another artificial ligand, GPI-GFP (Greco et al., 2001), which is anchored to the outer leaflet of the plasma membrane via a glycosylphosphatidylinositol (GPI) moiety (Paulick and Bertozzi, 2008), in the same setup as the soluble GFP and GFPext-CD8 (Figure 1F). A network of filopodial protrusions became apparent at the basal side of the epithelium (Figure 1F”, Supplementary Figure 1B), as with GFPext-CD8 expression. In addition, a lateral accumulation of signal was observed within approximately the first 4–5 cell diameters from the compartment boundary along the apical and lateral membranes. However, this ligand displayed a distinct behavior than GFPext-CD8: basal GFP protrusions are only observed emanating from the A compartment and immobilized on the basal surface of the P compartment, a behavior that may arise from differences in effective membrane tethering between GPI-anchored and transmembrane constructs (Hu et al., 2015) (see schemes in Supplementary Figure 2 for clarification).

**Figure 2.**
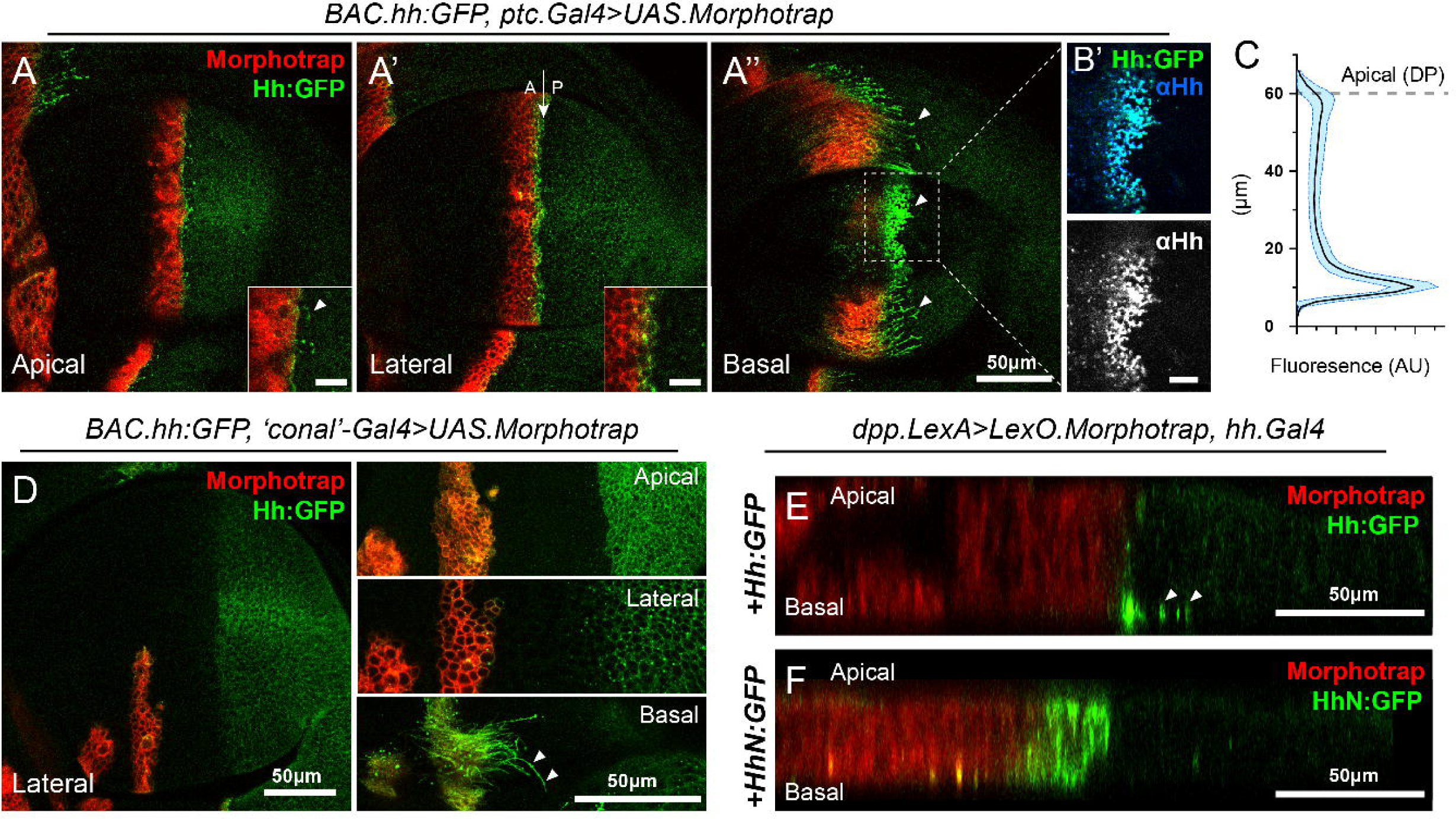
Hh immobilisation reveals Hh lipid-dependent association with cytonemes. A-A’’) Hh:GFP localization in apical, lateral and basal sections upon expression of Morphotrap in Hh receiving cells using the *ptc.Gal4* driver. Note that very short apical cytonemes are also observed (A and inset for magnification). In a lateral section, Hh accumulates on the membrane of the first two rows of cells immediately anterior of the A/P compartment boundary (inset for magnification) (A’). In the basal section, Hh is accumulated in Morphotrap expressing cytonemes that emanate from the A compartment cells (A’’). B’) Immunostaining using anti-Hh antibodies reveals accumulation in the same structures, indicating that Hh and GFP are accumulated together. C) Quantifications of the apical/basal Hh distribution after morphotrap expression in the Hh receiving cells. D) Clones expressing Morphotrap and situated far from the Hh source in a lateral view. Enlargement of the same clones in an apical, lateral and basal sections. Note that Hh accumulates preferentially in the basal part of the clone, but also in the region in between the clone and the Hh source. Note that Hh:GFP accumulation in the basal side of the clones is much higher than in the lateral or apical planes. Note also that cytonemes emanating from the clone show strong accumulation of Hh:GFP. Arrowheads indicate cytonemes. E) Apico-basal sections of a wing disc expressing Hh:GFP in P cells using *hh*.Gal4 driver and Morphotrap in A cells using *dpp*.LexA driver. The expression was limited to 48h by employing a *tub*.Gal80ts transgene. Most lipid modified Hh:GFP accumulates in the basal side (arrowheads). F) Same setup as in E but expressing HhN:GFP (non cholesteroylated form). In this case, Hh accumulates mainly in the apical side of the epithelium, with minimal basal and lateral accumulation.

### Hh immobilization reveals its lipid-dependent association to cytonemes

Having established the differential response of diffusible and membrane-bound ligands upon immobilisation, we examined the retention pattern of the morphogen Hh. For this purpose we employed a BAC containing the entire *hh* locus fused to GFP (Bac.*hh:GFP*) (Chen et al., 2017). Expression of morphotrap in the Hh-receiving cells using the *ptc*.Gal4 driver resulted in the basolateral accumulation of Hh:GFP in the two first rows of cells of the A compartment (Figure 2A’, A’’). Quantification of the GFP intensity along the apico-basal axis in these wing discs revealed that most Hh:GFP is immobilized on the basal side of the epithelia (Figure 2C). At this level, Hh:GFP locates on long protrusions emanating from the A compartment that invaded the P compartment (Figure 2A’’). Immunostaining using an anti-Hh antibody confirmed the immobilization of Hh protein on cytonemes (Figure 2B’). High resolution imaging confirmed the filopodial nature of these extensions, which also presented numerous thickenings (Supplementary Figure 3A). By contrast, in the apical and apico/lateral levels only one cell row accumulated Hh, with very short protrusions occasionally being observed (Figure 2A, A’ insets). These results suggest that Hh:GFP immobilization stabilizes cytonemes by stabilizing cell-cell interactions, where receiving cells expose a membrane tethered receptor, and the producing cells a membrane associated ligand.

**Figure 3.**
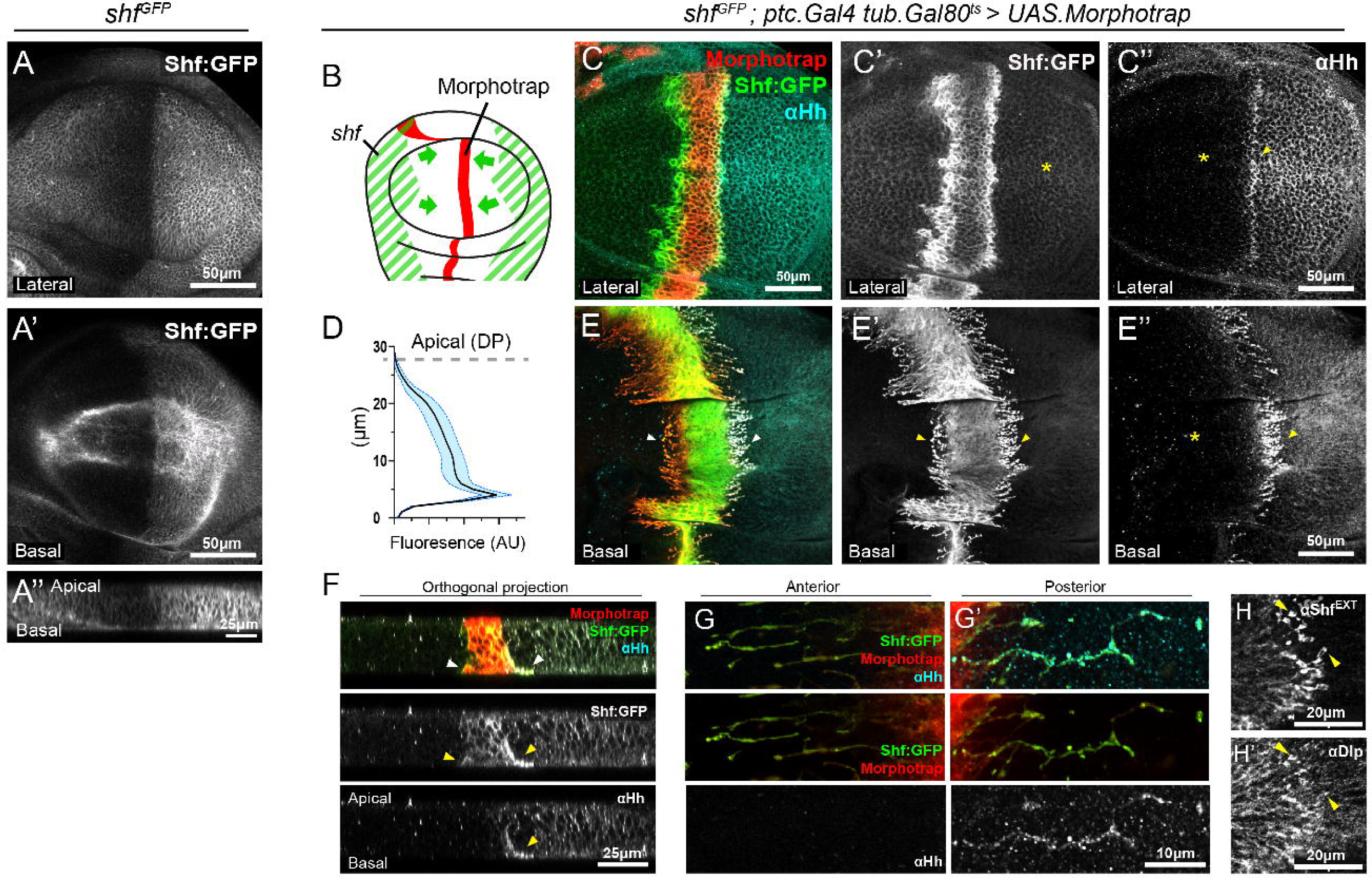
Shf:GFP is associated to membranes. A, A’) Expression pattern of Shf protein distribution in a *shf:GFP* wing disc in lateral (A) and basal sections (A’). B) Scheme representing the transcriptional pattern of *shf* gene in the wing disc, which is different from the Shf protein pattern distribution. Shf protein is stabilized in the P compartment. C-E’’) Wing imaginal disc showing the Shf and Hh protein distribution visualized by endogenous P-tagged Shf protein (Shf:GFP) and by the anti-Hh antibody after the expression of morphotrap for 24 hours under control of the *ptc.Gal4; tub.Gal80^ts^* driver. **D)** Quantification of apical versus basal Shf distribution after morphotrap expression in Hh-receiving cells. F) Orthogonal views showing the apico-basal distribution of Shf and Hh in the same disc. Yellow arrows indicate regions of colocalization. G–G’) High-magnification views of basal protrusions extending toward the anterior (G) and posterior (G’) compartments. Shf:GFP is detected in protrusions projecting in both directions, whereas Hh is preferentially enriched in protrusions oriented toward the posterior compartment. H, H’) Extracellular Shf (H) and Dlp (H’) detected with anti-Shf and anti-Dlp antibodies. Upon Shf:GFP capture by morphotrap, Hh and Dlp are co-recruited, predominantly in basal protrusions from A compartment cells that extend toward the P compartment (yellow arrows).

Cytonemes have been proposed to exchange morphogens in a synaptic-like process (González-Méndez et al., 2017; Huang et al., 2019). In fact, in the cytoneme-mediated transport model, the existence of direct cytoneme-cytoneme connection would be critical for these structures to deliver Hh to the farthest Hh-responding cells. We hypothesised that, if cytonemes are able to interact over long distances, expression of morphotrap in clones far from the border should also stabilize cytonemes. Supportingly, when morphotrap is expressed in clones induced in the A compartment, as far as 50μm away from the P compartment, Hh-GFP accumulates in basal protrusions (Figure 2D, inset). In these clones, cytonemes facing the P compartment were preserved after fixation, suggesting stabilization via cell-cell interactions with opposite projections originating from P compartment cells (Figure 2D, inset). When morphotrap-expressing clones were generated closer from the border the accumulation of Hh:GFP in these protrusions was much more pronounced (Supplementary Figure 3B).

To assess the dynamics of these protrusions in vivo, we performed live imaging in abdominal histoblasts, a system that allows robust visualization of thin cellular extensions and has been previously used to study cytoneme behavior (Bischoff et al., 2013). Live imaging of abdominal histoblasts expressing morphotrap under *ptc*.Gal4 revealed highly dynamic protrusions extending from morphotrap-positive cells (Supplementary Movie 1). In contrast, in histoblasts expressing Hh:GFP together with morphotrap under *ptc*.Gal4, protrusions formed a largely static network with little detectable movement (Supplementary Movie 2).

As described above, Hh lipid modifications are critical for gradient formation. Thus, we tested whether Hh association to cytonemes is dependent on these moieties. In particular, we investigated the retention pattern of both Hh:GFP and HhN:GFP, a version lacking the cholesterol modification (Callejo et al., 2006). We ectopically expressed either form using *hh*.Gal4, while simultaneously expressing morphotrap using *dpp*.LexA. The expression was limited to 48h by employing a *tub.*Gal80^ts^ transgene. Immobilization of wild-type Hh:GFP resulted in its accumulation in basal protrusions (Figure 2E), similar to those observed using the BAC.*hh:*GFP (Figure 2A’’ and Supplementary Figure 3A). In contrast, expression of the uncholesteroylated HhN:GFP did not stabilize cytonemes (Figure 2F). Instead, HhN:GFP accumulated in the first four rows of cells, labeling the plasma membranes along the apico-basal axis. In addition, uncholesteroylated HhN:GFP was also observed in larger puncta in the basolateral side of the disc (Figure 2F). The absence of uncholesteroylated Hh (HhN) in cytonemes has previously been reported in *Drosophila* wing imaginal disc (Callejo et al., 2011) and in vitro cultured cells (Bodeen et al., 2017).

Our results suggest that Hh is normally bound to the membranes of producing cells, that receiving cytonemes are exposed to large amounts of Hh, and that these processes are dependent on Hh lipids.

### Shf behaves as a membrane-associated protein

As mentioned earlier, the Shf transcript is present in the wing disc’s lateral regions (see Figure 3B). However, the Shf protein spreads extensively from its transcriptional domains(Glise et al., 2005; Gorfinkiel et al., 2005; Aguilar et al., 2024b), and contributes to the establishment of the Hh gradient along the A/P compartment boundary (see Figure 3A, A’). The high diffusivity of Shf, together with its role in the release of lipidated Hh from the P compartment (Bilioni et al., 2013; Glise et al., 2005; Gorfinkiel et al., 2005), led to the proposal that the Hh gradient could be formed by diffusion of the soluble Hh-Shf complex (Glise et al., 2005; Gorfinkiel et al., 2005). We previously generated a *shfGFP* knock-in allele, in which GFP is inserted into the endogenous *shf* locus (Figure 3A, A’) (Aguilar et al., 2024b), enabling direct assessment of Shf behaviour using the immobilization assay.

Expression of morphotrap in a stripe of cells of the A compartment using *ptc.*Gal4, led to the accumulation of Shf:GFP along the basolateral membranes, particularly on the two cell rows at both sides of the stripe (Figure 3C, C’ and quantified in D). This accumulation coincided with that of Hh only in the cells closest to the AP border, as revealed by anti-Hh immunostaining (Figure 3C’’). In the most basal side of the epithelia, Shf:GFP accumulated in cellular protrusions, which emanated from morphotrap-expressing cells oriented toward the laterals of the wing (Figure 3 E, E’, F). A similar stabilization behavior was observed in histoblasts carrying the endogenous shf:GFP allele together with morphotrap driven by ptc.Gal4 (Supplementary Movie 3). High-resolution wing disc views revealed filopodial extensions projecting toward both the anterior (A) and posterior (P) compartments (Figure 3G, G′). Notably, robust Hh accumulation was preferentially observed on cytonemes oriented toward the P compartment (Figure 3G′), whereas protrusions extending toward the A compartment displayed markedly reduced Hh retention (Figure 3G). We confirmed by anti-Shf immunostaining of non-permeabilized wing discs that the filopodial accumulation of Shf:GFP was extracellular (Figure 3H, see materials and methods). Interestingly, even though immobilization of Shf:GFP occurred at high levels not only on the most-basal filopodial extensions, but also elsewhere along the basolateral membranes (quantified in Figure 3D), Hh accumulated most prominently on basal cytonemes (Figure 3E’’, G and G’). In addition, the glypican Dally-like protein (Dlp), one of Hh coreceptors, was also found accumulated in stabilized cytonemes (Figure 3H’).

Considering that Shf is a diffusible protein, its localisation along stabilized cytonemes contrasts with the distribution when trapping soluble GFP (Figure 1 B), where GFP was homogeneously distributed along the membrane and does not stabilise cytonemes (Figure 1A, B). Conversely, Shf:GFP immobilization more closely resembles the capture of GPI-GFP (Figure 1C, D) and Hh:GFP itself (Figure 2E), suggesting that Shf is also anchored to membranes. Interestingly, these filopodia were stabilised both towards Hh-expressing and non-Hh-expressing cells (Figure 3 E, E’), arguing against Hh-dependent membrane tethering of Shf. Thus, we hypothesised that a membrane-bound factor expressed ubiquitously in wing discs must anchor Shf to the basolateral membrane.

### Ihog stabilises Shf at cell membranes via its Ig domains

The Hh co-receptor Ihog is a type I transmembrane protein. It contains two fibronectin type III domains and four Ig domains in their extracellular region. Previous studies have established that Ihog, as well as its mammalian paralog CDON, serve as interaction “hubs” for most of the extracellular components of Hh pathway (Bilioni et al., 2013; McLellan et al., 2006; Simon et al., 2021; Wierbowski et al., 2020; Wu et al., 2019; Yao et al., 2006). In the wing disc, Ihog is involved in the maintenance of Shf levels and when overexpressed, Shf extracellular levels are increased (Avanesov and Blair, 2013; Bilioni et al., 2013) specially at cytonemes (Bilioni et al., 2013). We thus tested whether Ihog interaction with Shf could also be direct. Structure modeling of Ihog using AlphaFold 3 (Abramson et al., 2024) predicted with high confidence that the four Ig domains adopt a “closed” conformation, with high model confidence across the Ig-Ig interfaces (Supplementary Figure 4A). This feature was conserved for the structure of Boi (Supplementary Figure 4 B, B’). Simultaneous modeling of Ihog and Shf revealed a putative interaction, formed by the EGF domains of Shf and the Ig1/Ig4 of Ihog (Figure 4A, B, and C). Notably, the loss-of-function allele *shf²* mutation maps to one of the EGF domains (Glise et al., 2005; Gorfinkiel et al., 2005) predicted to contact Ihog (Figure 4C), providing independent genetic support for this interface. This interaction was also preserved when both Boi and Shf were modelled together (Supplementary Figure 4B, B’).

**Figure 4.**
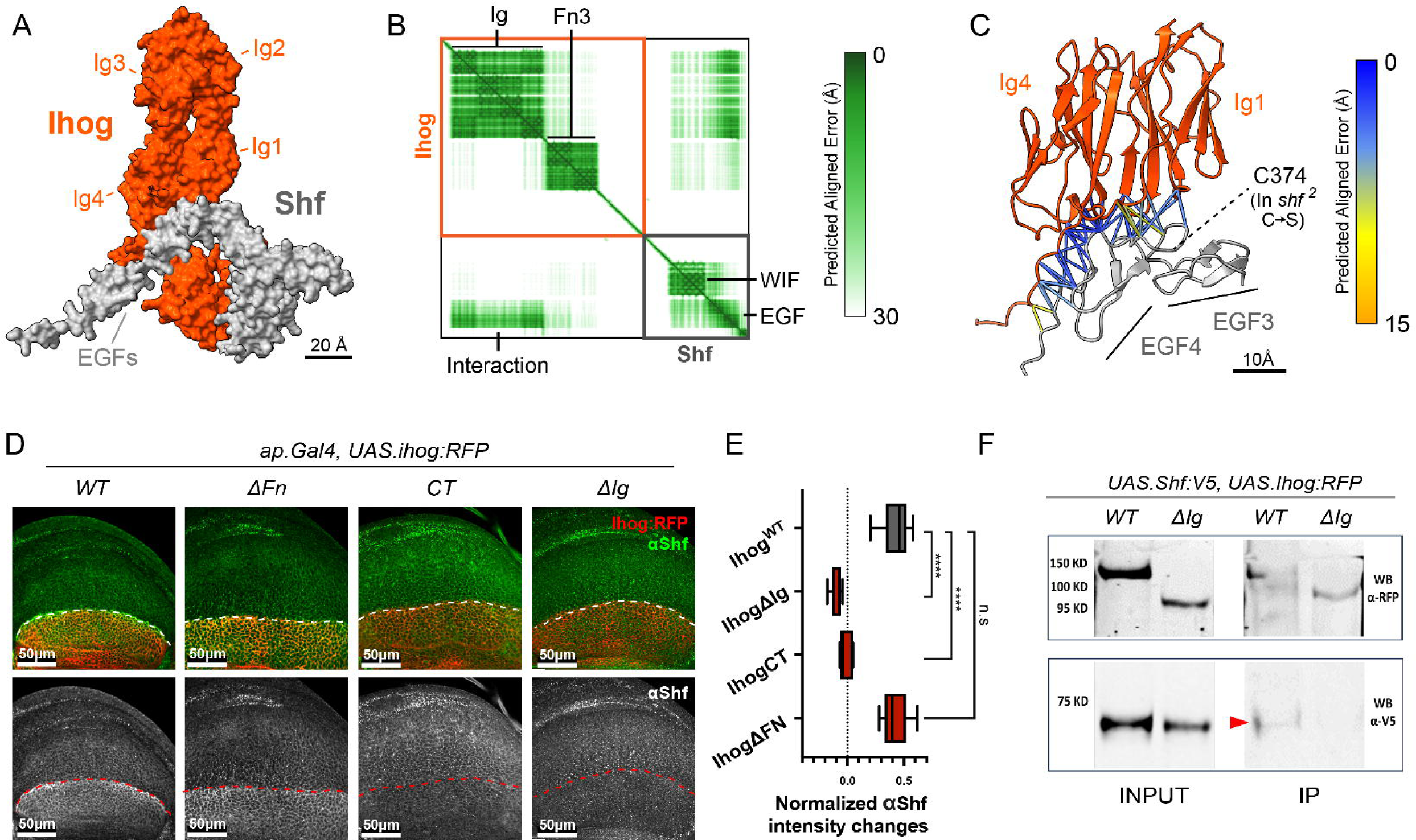
The Ig domains of Ihog interact with Shf. A) AlphaFold 3 model of Ihog and Shf. Ihog Ig domains (indicated) adopt a folded conformation in which Ig1-Ig4 and Ig3-Ig2 interact. This folded structure forms the surface with which Ihog interacts with Shf. Only globular domains are shown in the structural model. B) AlphaFold confidence matrix indicating the predicted aligned error of the model in A. Modeling was performed using full-length proteins. C) Detail of the interaction between Ig domains of Ihog and the EGF domains of Shf. In Blue-Orange Palette, the confidence of the interaction is measured using ChimeraX; the shf² mutation site is indicated within the interacting EGF domain.D) Shf protein accumulation (visualised with anti-Shf antibody) induced by overexpression of different Ihog mutant forms in the dorsal compartment of wing discs using the *ap.Gal4; tub.Gal80^ts^* driver, keeping the ventral compartment as an endogenous control: *UAS.Ihog:RFP*; *UAS.Ihog*Δ*Fn:RFP; UAS.IhogCT:RFP*; *UAS.Ihog*Δ*Ig:RFP*. Note that the expression of Ihog form lacking the Ig domains is unable to recruit Shf. This is not the case for the Ihog form lacking the FN domains. E) Quantifications of Shf accumulation in the dorsal compartment by the expression of the Ihog mutant forms relative to the endogenous control (ventral compartment). (F) Western blots after co-immunoprecipitation assay in salivary gland protein extract expressing Shf:V5 (53 kDa) and Ihog:RFP (135 kDa) and IhogΔIg:RFP (95 kDa) proteins using an anti-RFP antibody. The upper panel shows the immunoprecipitation of Ihog:RFP and IhogΔIg:RFP proteins visualised with an anti-RFP antibody. The lower panel shows the co-immunoprecipitation of WT Shf (Shf:V5) protein visualised with anti-V5 antibody. Note that Shf does not co-immunoprecipitate with the Ihog protein forms lacking the immunoglobulin domains.

To further analyse the Ihog/Shf interaction *in vivo*, we overexpressed different Ihog mutant forms in the wing disc and analysed the Shf recruitment using a specific anti-Shf antibody (Figure 4 D, E). Unlike wild-type Ihog:RFP, the expression of either IhogCT:RFP (containing only the intracellular C-terminal part of Ihog) or IhogΔIg:RFP failed to accumulate Shf (Figure 4D, E for quantifications), indicating that Ihog-Shf interaction occurs via the extracellular part of Ihog and specifically through the Ig domains. We further confirmed these results by a co-immunoprecipitation analysis, coexpressing the different Ihog mutant forms tagged with RFP and Shf:V5 in salivary glands. The IhogΔIg mutant failed to co-immunoprecipitate with Shf, confirming the requirement of the Ig domains for the Ihog/Shf interaction (Figure 4F).

### Ihog Ig domains play distinct roles on Hh stabilization and signal transduction

Shf appears to play a dual role in Hh-producing cells: it stabilizes Hh at the cell surface (Avanesov et al., 2012; Bilioni et al., 2013) and facilitates its release to form a gradient (Glise et al., 2005; Gorfinkiel et al., 2005; Bilioni et al., 2013). The Ihog fibronectin 1 (Fn1) domain has been implicated in direct interaction with Hh (McLellan et al., 2006; Yao et al., 2006; Bilioni et al., 2013; Simon et al., 2021; Yang et al., 2021). Nevertheless, we observed that IhogΔIg:RFP, which preserves intact FN domains, failed to stabilize Hh (Figure 5B, B’). This result suggests that Shf cooperates with Ihog to retain Hh via the Ihog Ig domains, potentially forming a tripartite Ihog-Shf-Hh complex.

**Figure 5.**
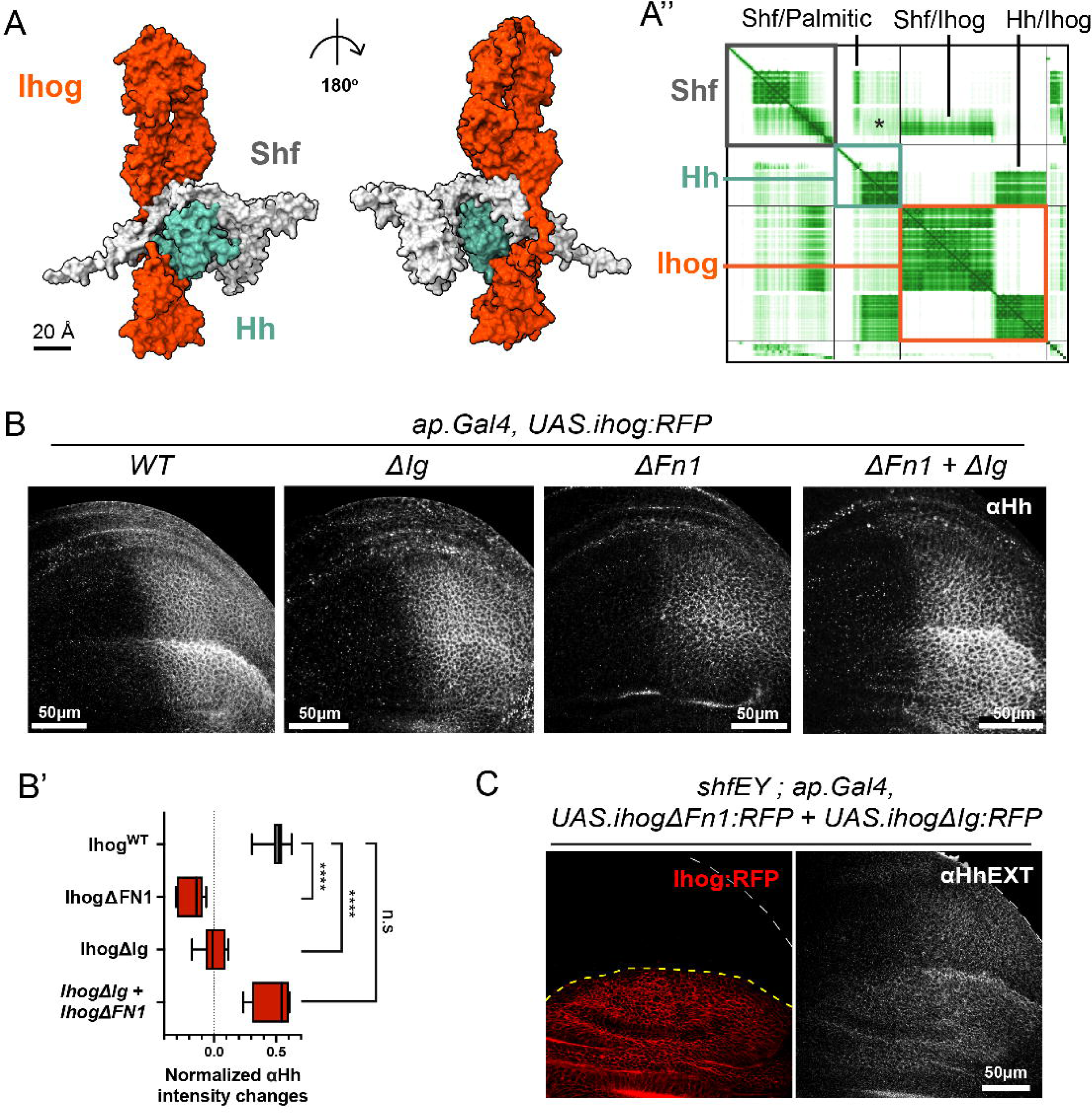
Shf is required for Ihog-mediated stabilization of Hh. **A)** Structural prediction of the Ihog–Hh–Shf tripartite complex. **A’)** AlphaFold confidence matrix showing predicted aligned error for the model in (A). Each protein is color-coded; interactions are indicated in black. Shf–Hh interaction is restricted to residues surrounding the palmitoylated region of Hh and the lipid moiety. Only globular domains are displayed in the surface model. Predictions were computed using full-length Hh and Shf and the globular domains of Ihog. **B)** Hh accumulation (anti-Hh) in the dorsal compartment of ap.Gal4; tub.Gal80ts wing discs after overexpression of Ihog variants: UAS.Ihog:RFP (WT), UAS.IhogΔIg:RFP, UAS.IhogΔFn1:RFP, and the combined mutant UAS.IhogΔIg:RFP/UAS.IhogΔFn1:RFP. Hh accumulation is restored when one construct retains the Ig domains and the other the Fn1 domain. **B’)** Quantification of dorsal Hh accumulation relative to the endogenous ventral control (see Materials and Methods). **C)** Extracellular Hh staining in discs overexpressing UAS.IhogΔIg:RFP together with UAS.IhogΔFn1:RFP using ap.Gal4; tub.Gal80ts.

Modeling of this putative complex, confirmed the feasibility of such an arrangement (Figure 5A). In the model, Shf engages the palmitic moiety of Hh through the pocket in its WIF domain while simultaneously contacting the Ig domains of Ihog via its EGF repeats (contacts described in Figure 4). Consistent with this interface, the *shf²* allele, which phenocopies *shf* null mutants, harbors a C374S substitution within the EGF repeats region (Glise et al., 2005; Gorfinkiel et al., 2005) predicted to contact Ihog. In parallel, Ihog and Hh maintain the previously described interaction through the Fn1 domain (McLellan et al., 2006) (Figure 5A and A’). We detected no direct contacts between the EGF domains of Shf and Hh in this model (Figure 5A’, asterix).

Importantly, an Ihog variant carrying three substitutions in the Fn1 domain that disrupt Hh binding (IhogFn1***) (Supplementary Figure 5A) still accumulated Shf on cytonemes, similarly to wild-type Ihog (Figure 5B), indicating that the interaction between Ihog and Shf is independent of Hh. Consistent with this, a form of Ihog without the FN domains (IhogΔFN:RFP), unable to interact with both Hh and glypicans (Simon et al., 2021; Yang et al., 2021), retained the ability to recruit Shf (Figure 4D). Further demonstrating the dispensability of Hh and glypicans on the interaction between Ihog and Shf and supporting a direct interaction between Shf and Ihog.

To control for potential interference from endogenous Ihog, we examined endogenous Ihog levels upon expression of Ihog:RFP and its mutant derivatives. We observed that ectopic expression of all Ihog forms reduced the Ihog–GFP levels from a genomic BAC.Ihog-GFP transgene encompassing the endogenous ihog regulatory and coding sequences (see Supplementary Figure 6), indicating that the phenotypes observed reflect the specific properties of each mutant construct.Ihog dimerization through the Fn1 domain has been shown to mediate the high-affinity interaction between Ihog and Hh (McLellan et al., 2006). Unexpectedly, co-expression of complementary Ihog mutants (IhogΔIg:RFP / IhogΔFn1:RFP or IhogΔIg:RFP / IhogΔFN:RFP) restored Hh accumulation (Figure 5B, B′; Supplementary Figure 5), although none of these Ihog forms can independently accumulate Hh (Figure 5B, B’ and Supplementary Figure 5). Because one partner in each pair lacks the Fn1 homodimerization domain, this rescue is unlikely to rely on canonical Ihog homodimerization. (Yang et al., 2021). Besides, co-expression of Ihog mutants that are not able to interact with Hh, such as IhogFn1***:RFP and IhogΔFN:RFP, did not rescue the ability of Ihog to accumulate Hh (Supplementary Figure 5) (Simon et al., 2021; Wu et al., 2019; Yang et al., 2021).

**Figure 6.**
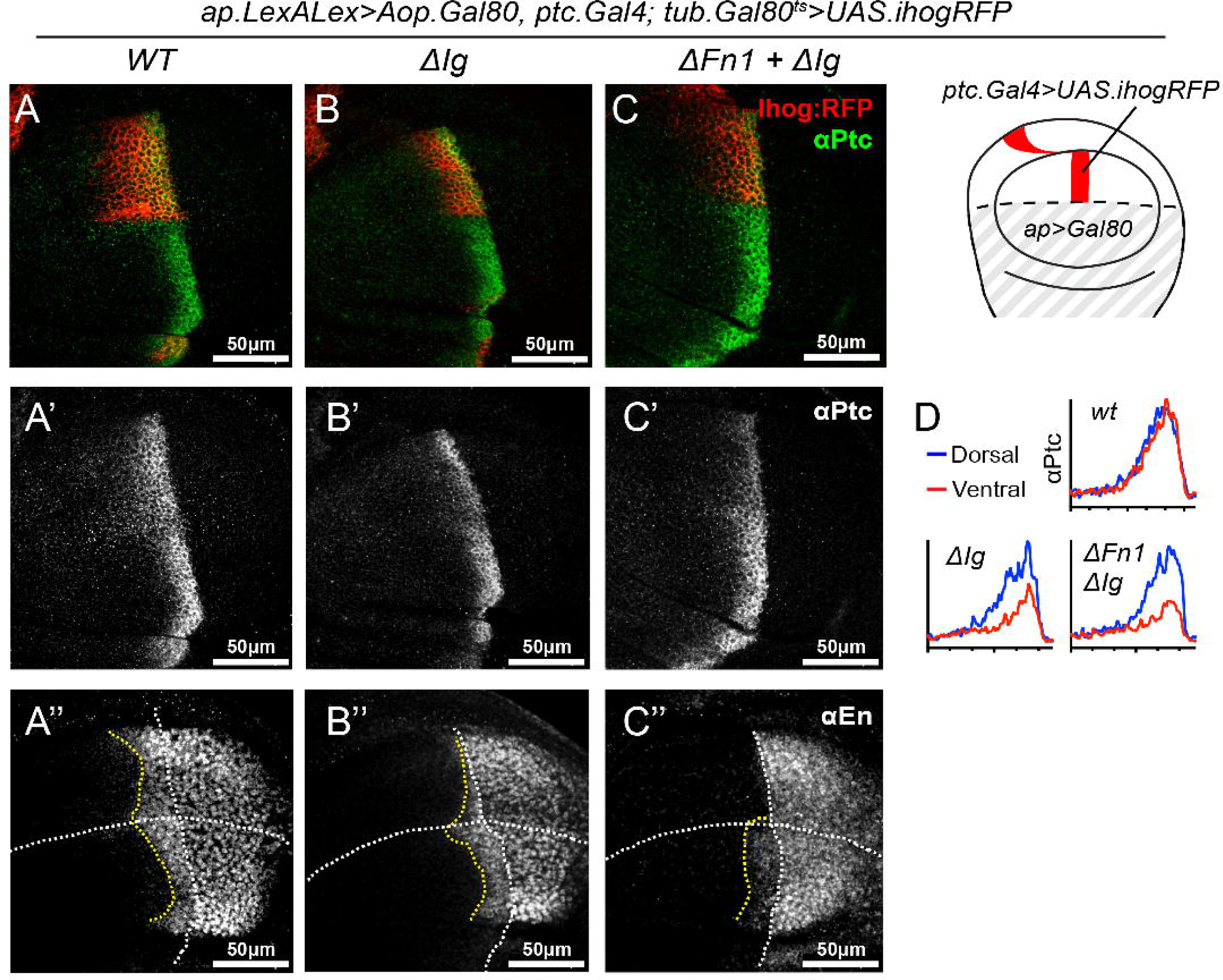
Ihog Ig and Fn1 domains must act in cis to support Hh co-receptor function. A–C’’) Expression of high-threshold Hh targets Ptc and En in wing discs after overexpression of Ihog variants using LexAop.Gal80; ap.LexA; tub.Gal80ts. The dorsal/anterior domain serves as the experimental region with the ventral domain as internal control (schematic shown). Genotypes include UAS.Ihog:RFP (WT), UAS.ral (red) compartments for the indicated Ihog variants (see Materials and Methods).IhogΔIg:RFP, and the mutant combination UAS.IhogΔIg:RFP/UAS.IhogΔFn1:RFP. Expression of UAS.IhogΔIg:RFP strongly reduces target gene activation. Although the double-mutant combination restores Hh stabilization in producing cells (Figure 5), Ihog co-receptor function is not rescued in receiving cells. D) Quantification of Ptc levels in dorsal (blue) versus ventral.

In addition to its role in stabilising Hh at the plasma membrane (Bilioni et al., 2013; Callejo et al., 2011; Yan et al., 2010), Ihog has also been described as an Hh coreceptor (Simon et al., 2021; Yang et al., 2021; Zheng et al., 2010). We tested the possible involvement of the Ig domains in Hh reception by evaluating the effect that the expression of the different Ihog mutants and mutant combinations had in Hh-receiving cells (Figure 6A-D’’). We drove the expression of the wild type and the different Ihog mutant forms exclusively in the ventral area of the A compartment of the wing disc, leaving the dorsal/anterior area unaffected (see scheme of the Gal4 induction at the right hand side of Figure 6). Expression of wild-type Ihog:RFP resulted in the flattening of the signalling gradient. Conversely expression of the Ihog construct lacking the Ig domains (IhogΔIg:RFP) resulted in the dominant reduction of signaling (Figure 5B-B’’), similar to that observed by expressing Ihog form mutated in the Fn1 domain (Simon et al., 2021) (Supplementary Figure 7C). In those lines, all the pairwise combinations of Ihog mutant proteins that were able to restore Hh accumulation in the P compartment (Figure 4B) reduced Hh signalling when expressed in the A compartment (Figure 6C-C’’ and Supplementary Figure 7C, quantified in Figure 6D). This suggests a different behaviour of Ihog-Shf-Hh complex during Hh stabilization and signal transduction.

**Figure 7.**
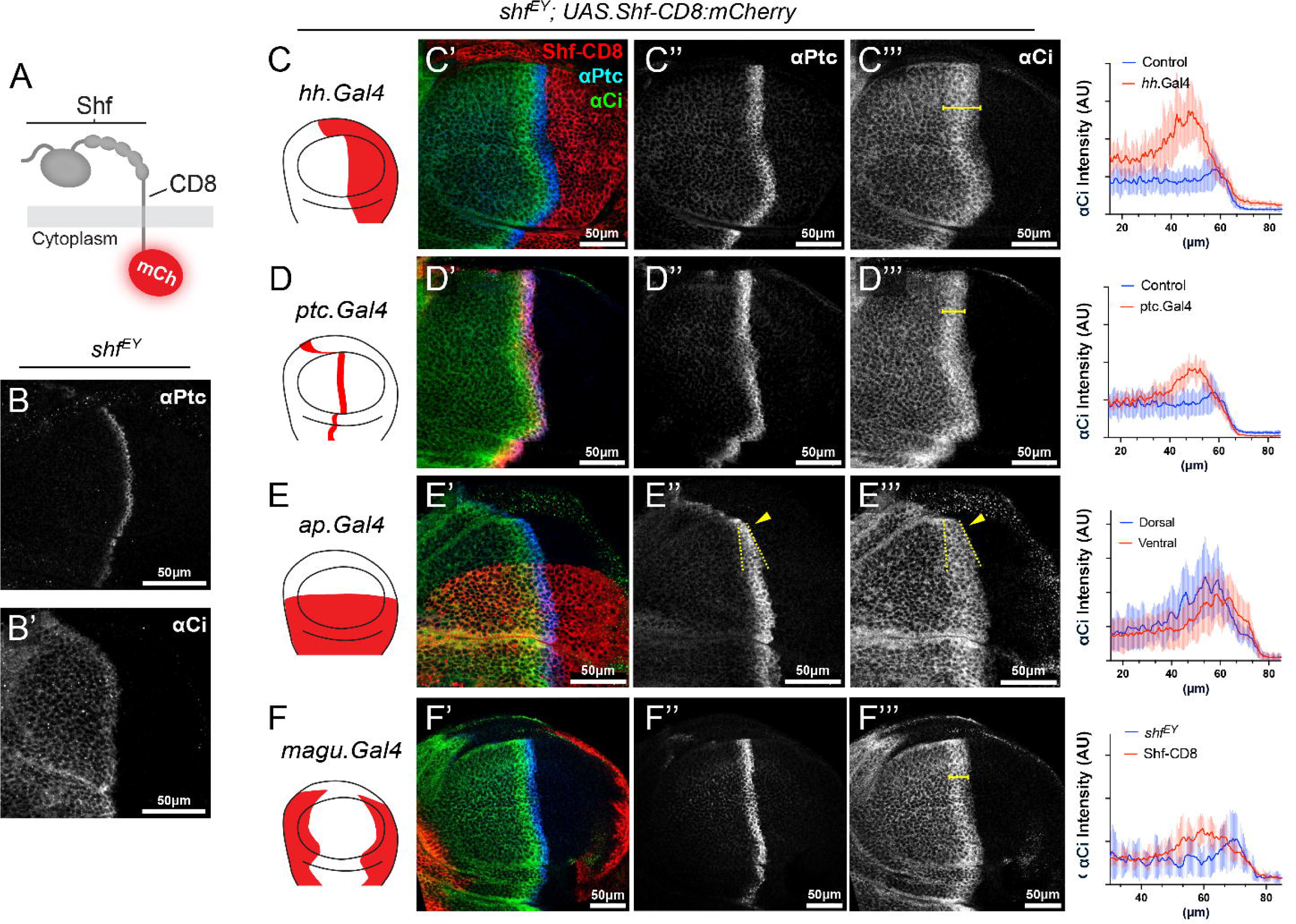
Membrane-anchored Shf protein is able to rescue Hh signalling in *shf^EY^* null mutants. A) A drawing of a form of Shf attached to membranes by CD8 sequences and intracellularly tagged to mCherry (Shf-CD8:mCh). B, B’) *shf^EY^*wing disc stained with anti Ptc (B) and anti Ci (B’) antibodies as Hh signalling reporters. Note the absence of the Hh gradient. C-F’’’) Effects on Hh signalling after expression of *UAS*.Shf-CD8:Ch using the *hh*.Gal4 (C), *ptc*.Gal4 (D), *ap*.Gal4 (E) and *magu*.Gal4 (F) drivers in a *shf^EY^*null mutant wing imaginal disc. The effects on Hh signalling is reported by the expression of Ptc and Ci using specific antibodies. Note that the expression of the Shf-CD8:mCh is able to rescue Hh signalling when expressed in the A compartment (D’, D’’, D’’) and in a non-autonomous manner when it is expressed in the P compartment (C’, C’’, C’’’). In the case of expressing Shf-CD8 in the dorsal compartment of the wing disc using *ap*.Gal4 line, there is also a non-autonomous rescue of Hh signalling in a gradient-like shape in the ventral compartment (E‘, E’’, E’’’, yellow dotted lines and arrows). Note also that when Shf-CD8:mCh is expressed distant from the Hh reception area using the *magu*.Gal4 driver (red) there is also a non-autonomous rescue of Hh signaling (F’, F’’, F’’’).

Most extracellular pathway components are present both in producing and receiving cells, Ptc being the main asymmetrically distributed protein, responsible for triggering the signaling response in receiving cells. Thus, we modeled the putative complex formed by Shf-Hh-Ihog-Ptc using Alphafold 3 (Supplementary Figure 8A).The model supports the simultaneous interaction of Ptc, Ihog, and Hh, supported by previous experimental studies (Zheng et al., 2010), while remaining compatible with Ihog–Shf binding through the Ig domains. Notably, the model predicts that engagement of Ptc disrupts the Shf WIF–palmitate interaction with Hh (Supplementary Figure 8B, asterisk).

**Figure 8.**
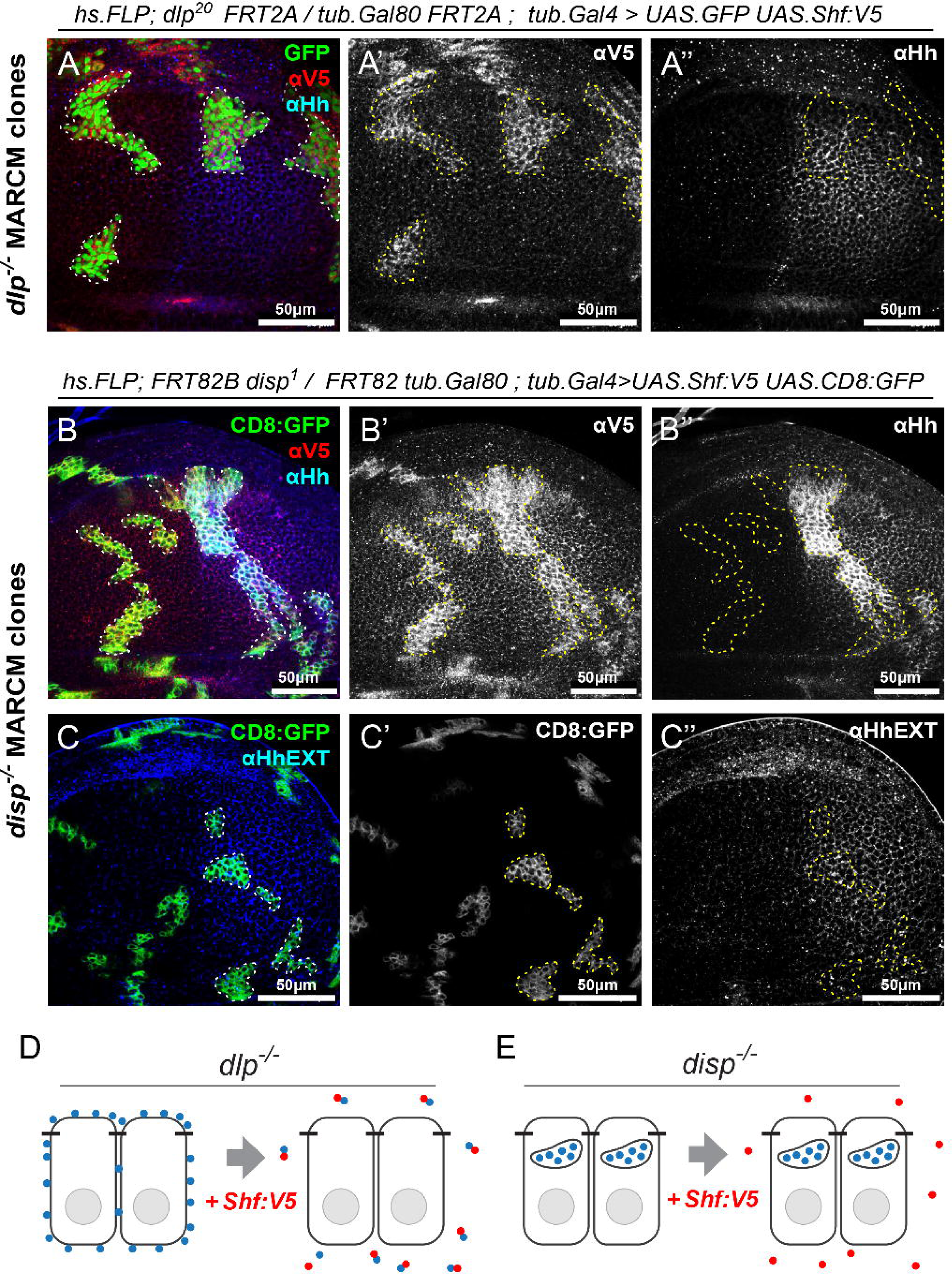
Shf rescues Hh accumulation in *dlp^-/-^* cells but not in *disp^-/-^* cells. (A-B) *dlp^-/-^* (A) and *disp^-/-^* (B) mutant clones labeled by the expression a nuclear GFP (green) (A) and *UAS*.CD8:GFP (green) (B) respectively while overexpressing a *UAS*.shf:V5 in the same cells. For both Hh and Shf:V5 specific anti-Hh and anti-V5 antibodies were used. It can be observed that the overexpression of a Shf:V5 construct rescues Hh accumulation in *dlp*^-/-^ clones (A’’), whereas this rescue does not occur in *disp*^-/-^ mutant cells (B’’). (C) A wing disc containing *disp^-/-^* mutant clones labeled by the expression of a *UAS*.CD8:GFP in which only the extracellular Hh protein is labelled (C’’). The absence of the Hh ex-vivo staining indicates that Hh accumulation in *disp^-/-^* cells is intracellular (C’’).

### Membrane-bound Shf allows long distance Hh signalling non-autonomously

Shf appears to facilitate the release of Hh from the surface of producing cells, consistent with a function in solubilising Hh to form long-range gradients by diffusion (Glise et al., 2005; Gorfinkiel et al., 2005). Nevertheless, here we have also shown the association of Shf to plasma membranes and to cytonemes (Figure 3E, E’, G, H; Supplementary Figure 7A and B). We hypothesised that if Hh travels anchored to cytoneme membranes, Shf function should not be dependent on its diffusion, as long as it is present at the sites of contact between cells. To test this hypothesis we constructed a membrane bound Shf by fusing Shf to the extracellular region of CD8:mCh.

First, to analyse the spreading ability of Shf-CD8 and wild type Shf (Shf-V5), we expressed either of them in the peripodial membrane using the *ubx*.Gal4 driver (see scheme in Supplementary Figure 9A) in a *shf^EY^*mutant background. Wild-type Shf was able to restore Hh signaling in the disc proper (Supplementary Figure 9C, C’, and C’’ for *shf^EY^*rescue compared with *shf^EY^*mutant disc and adult wing in Supplementary Figure 9B-B’’). No rescue of Hh signalling was observed when Shf-CD8 was expressed in this setup (Supplementary Figure 9D, D’, D’’), which demonstrates that Shf-CD8 is unable to spread between tissues because it is not cleaved from its membrane tether.

**Figure 9.**
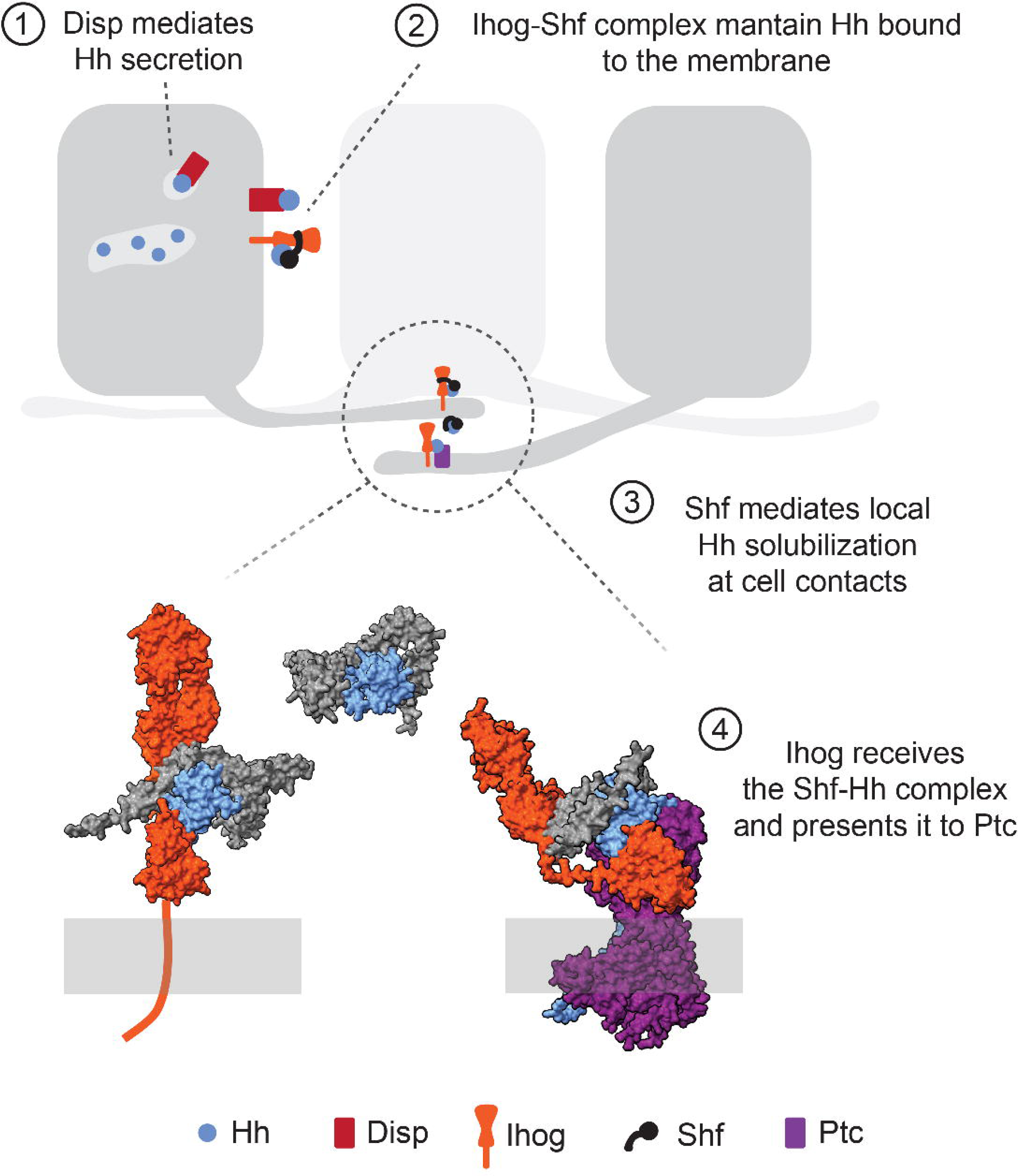
Model of the critical steps of Hh transport proposed in this study. Step 1: Disp mediates the secretion of Hh and may be involved in its intracellular trafficking from the apical to the basal side of the wing disc epithelium. Step 2: Ihog and Shf work together to maintain extracellular Hh bound to the membranes of Hh-producing cells. Step 3: Shf mediates local solubilisation at cytoneme-cytonemecontacts. Step 4: Ihog receives the Shf-Hh complex and transfers Hh to Ptc via direct interaction with Shf through the Ig domains, and with Ptc and Hh through the FN domains.

We then analysed whether Shf-CD8 was able to restore Hh signaling in *shf^EY^* mutants (Figure 7 B,B’) when expressed in either the A or in the P compartments of the disc proper (see schemes in Figure 7C and D). Expression of Shf-CD8 in the P compartment using *hh*.Gal4 (Figure 7C) resulted in the restoration of Hh signalling, as revealed by anti-Ptc and anti-Ci immunostaining (Figure 7C’-C’’’ and quantifications). A considerable rescue, if not as good as that observed using *hh*.Gal4, was obtained when Shf-CD8 was expressed in the A compartment using *ptc*.Gal4 (Figure 7D-D’’’ and quantifications). These results suggest that the presence of Shf in either producing or presenting membranes is enough to facilitate Hh signalling, and that Shf diffusion is dispensable for Hh gradient formation.

To test the non-autonomous activity of Shf-CD8, we drove its expression using *ap*.Gal4 in *shf^EY^* mutants. In this setup, Hh signalling gradient was restored in the dorsal compartment (Figure 7E), whereas a graded restoration was observed in the ventral compartment. The lowest Hh-target levels were activated in cells furthest from the DV boundary (Figure 7E’-E’’’ and quantifications).

To confirm the non-autonomous ability of Shf-CD8 to rescue Hh signalling in *shf^EY^* mutants, we expressed Shf-CD8 far away from the A/P compartment boundary under the control of the *magu*.Gal4 driver in a *shf^EY^* mutant background (Figure 7F). Although *magu* expression is located at the most distal ends of the A and P compartments (Figure 7F and F’ red channel, Supplementary Figure 10A), membrane-anchored Shf was able to partially rescue Hh signalling (Figure 7F’-F’’’ and Supplementary Figure 10B-B’’ compared with Supplementary Figure 10D,D’). Finally, in order to test whether cytonemes are responsible for the non-autonomous rescue of Shf-CD8, we co-expressed *scar.RNAi,* an RNAi that has been shown to disrupt filopodia (Bischoff et al., 2013), together with Shf-CD8, again using the *magu*.Gal4 driver in a *shf^EY^* mutant background. This set-up resulted in a reduction in the Hh signalling rescue (Supplementary Figure 10C-C’’). Together, these results suggest that Shf-CD8-loaded cytonemes can still reach the signalling zone at the A/P compartment boundary, which in turn facilitates Hh presentation to recipient cells.

### Shf facilitates the release of Hh from the surface of producing cells

In addition to its role stabilizing Hh at the cell surface, Shf also facilitates Hh release from P compartment cells to form a gradient in the A compartment (Glise et al., 2005; Gorfinkiel et al., 2005). We hypothesised that, to mediate Hh release, Shf must be able to interact with Hh independently of Ihog, which remains associated with the P compartment cells. Thus, we modeled the interaction of Shf and Hh in the absence of Ihog. Remarkably, Shf is predicted to interact directly with Hh via its EGF repeats (Supplementary Figure 11A-B). This interaction was never predicted when Ihog is present (Figure 5A, asterisk). Moreover, the Shf EGF repeat/Hh interaction was compatible with that of the Shf WIF domain/Hh palmitic acid (Supplementary Figure 11A’). This hypothesis is supported by the fact that Shf interacts with Hh through both the WIF domain and the EGF repeats. The *shf^919^* mutant allele, which behaves as a *shf* null, has a point mutation (C638S) in one of the EGF repeats predicted to interact with Hh (Supplementary Figure 11B). This model suggests that Shf might be able to exist both in a membrane associated form, when linked to Ihog and Hh, or in a soluble form, linked to Hh.

Motivated by the structural predictions, we tested whether Shf overexpression can change the stoichiometry of the complex, releasing Hh from cells in which it has been ectopically accumulated by Ihog overexpression. As a consequence of the Hh retention by Ihog (Supplementary Figure 12F-F’’), Hh signaling is compromised, monitored by the expression of Ci and Ptc (Yan et al., 2010; Bilioni et al., 2013, Avasenov and Blair 2013) (Figure Supplementary 12A-A’’). This effect on Hh retention (Supplementary Figure 12F-G’’) and signaling is counteracted by the coexpression of both Ihog and wild type Shf or Shf-CD8 (Figure Supplementary 12B, C’’).

Other mutant conditions where Hh is accumulated are in Dispatched (Disp) and the glypican Dlp mutant cells which both have also been shown to participate in Hh release from producing cells; Hh being accumulated inside both *disp^-/-^* (Burke et al., 1999; Callejo et al., 2011) (Supplementary Figure 13B) and *dlp^-/-^* (Avanesov and Blair, 2013; Callejo et al., 2011) (Supplementary Figure 13A) null clones causing a reduction of the Hh signalling. Yet, the sequence of action of the three components remains unknown. We first tested whether Shf could mobilize Hh in *dlp* null clones, in which the accumulation of Hh is extracellular (Avanesov and Blair, 2013). To that end, we generated *dlp* null clones in which we simultaneously expressed wild type Shf (*UAS*.Shf:V5) using the MARCM technology. Anti-Hh immunostaining revealed that Hh did not accumulate extracellularly, in contrast to *dlp^-^*^/-^ clones alone (Figure 8A-A’’). As for Dlp, we induced MARCM *disp^-/-^* clones that overexpressed *UAS*.Shf:V5 (Figure 8B). Unlike in *dlp^-/-^*mutant cells, Shf was not able to rescue Hh accumulation (Figure 8B’’) suggesting that in absence of Disp, Shf is unable to release Hh from the cell surface. Indeed, extracellular Hh does not accumulate in *disp^-/-^* clones, demonstrating that Hh was accumulated intracellularly in the absence of Disp (Figure 8B, C’’). These results suggest that Shf is only able to interact with Hh once it has reached outer surface of the plasma membrane, as it is the case in *dlp* mutant clones (see scheme in Figure 8D), but not in the case when it is still intracellular, as it is the case in *disp* clones (see scheme in Figure 8E).

Finally, to further clarify the order of action of the pathway components involved in Hh release, we carried out a functional genetic epistasis using RNAis against Disp, Ihog, and Dlp, and visualised their effect by Hh protein levels using the anti-Hh antibody. Using this approach, we observed that the effect of Disp RNAi prevailed over that of Ihog RNAi (Supplementary Figure 14), suggesting that Ihog’s role on Hh stabilization requires the prior action of Disp. On the other hand, the effect of Ihog RNAi prevailed over that of Dlp RNAi (Supplementary Figure 14).

Together, these results suggest that Hh is first secreted by Disp, that Ihog retains its in the membrane only once it has been secreted, and that Shf and Dlp mediate its release from the cell surface.

## Discussion

### Cytoneme-mediated Hh gradient formation

Two main models of Hh dispersal have been proposed to explain the underlying mechanism of Hh gradient formation in the wing imaginal disc: the diffusion model and the cytoneme model (reviewed in: González-Méndez et al., 2019). The diffusion model proposes that Hh is released through the apical side of the epithelium, where it would diffuse as oligomers or exovesicles to form a signaling gradient (Ayers et al., 2012, 2010; D’Angelo et al., 2015; Gallet et al., 2006; Hurbain et al., 2022). In contrast, the cytoneme model proposes that Hh forms the long-range gradient in the basal region of the epithelia by means of cytoneme contacts (Bilioni et al., 2013; Bischoff et al., 2013; Callejo et al., 2011; Chen et al., 2017; González-Méndez et al., 2017; Gradilla et al., 2014; Simon et al., 2021). Exosomes associated with cytonemes have also been shown to carry Hh. MVBs loaded with Hh move along cytonemes and are released as exosomes at their contact sites (Gradilla et al., 2014). The contact sites between cytonemes of the P and A compartments could facilitate the reception and transfer of Hh (Chen et al., 2017; González-Méndez et al., 2017; 2020). However, the mechanism of Hh exchange between producing and receiving cells remains unclear, and the relative contribution of these models has been difficult to resolve, in part because of the difficulties in manipulating the subcellular localisation of Hh using genetic tools, and also because of the rapid dynamics of Hh secretion, distribution and internalisation.

In this study, we have revisited the question of Hh dispersal by using protein binders (Matsuda et al., 2022; Schnider et al., 2024). This approach allows direct in vivo interrogation of extracellular ligand behavior with spatial precision. To date, the trapping of extracellular ligands on the surface of receiving cells had been tested for BMP (Bauer et al., 2023; Harmansa et al., 2017, 2015; Matsuda et al., 2022, 2021) and Wnt (McGough et al., 2020; Mii et al., 2021; Pani and Goldstein, 2018). In these studies, the immobilized ligands resulted in their accumulation along the membrane, similar to the trapping of secreted GFP we report here (Figure 1). On the contrary, immobilization of Hh:GFP resulted in the accumulation of the ligand in the most basal side of the disc (Figure 2), suggesting a biased secretion towards this side of the epithelia. Most strikingly, immobilization also resulted in the stabilization of receiving cytonemes, which extended towards the source of the ligand (Figure 2; Supplementary Figure 3).

These differences with other morphogens could suggest an intrinsically different extracellular behaviour of Hh. In fact, the retention pattern of Hh:GFP more closely resembles that of membrane-bound GFP or GPI:GFP (Figure 1). Yet, these three conditions also present slight differences. In the case of Hh GPI:GFP, cytonemes are only stabilized towards Hh producing cells (Figure 2), while the trapping of membrane-bound GFP also stabilizes the contact in the other direction (Figure 1). We interpret these differences as reflecting the intermediate effective membrane affinity of lipid-modified Hh compared with the strongly membrane-tethered GFP:CD8 construct. In those lines, filopodia appear to fragment when cellular contacts are constantly stabilized by the morphotrap/GFP:CD8 bond (Figure 2E), probably due to the immobilization of these structures by morphotrap.

In addition, we demonstrated that the trapping of Hh is dependent on its lipids modifications (Figure 2). These lipid moieties have been shown to link Hh to membranes (Porter et al., 1996), which further favours the interpretation that the immobilization assay is revealing the high affinity of lipid-modified Hh for the extracellular membranes. Wnt ligands are also lipid modified (Willert et al., 2003), however, no evidence of filopodial stabilization has been reported in the previous studies using membrane trapping (McGough et al., 2020; Mii et al., 2021; Pani and Goldstein, 2018). In contrast, other experimental approaches have described specific Wnt transport by cytonemes in Drosophila (Hatori et al., 2021) and in vertebrates (Mattes et al., 2018; Mittermeier and Virshup, 2022; Routledge et al., 2022; Routledge and Scholpp, 2019; Stanganello and Scholpp, 2016; Waghmare and Page-McCaw, 2021; Zhang et al., 2024).

Here, we report that the immobilisation of the highly diffusible GFP:Shf molecule using the GFP morphotrap exhibits behaviour that is very similar to that of Hh:GFP (Figure 3). Thus, despite being biochemically diffusible, Shf behaves in vivo as a membrane-associated component of the cytoneme transport machinery. Since Shf is a factor specific for the Hh signalling pathway in Drosophila, its colocalization with Hh in cytonemes strongly suggests that the Hh gradient is formed using cytonemes. The colocalization of Shf and Hh at the cytoneme contact sites indicates a dispensable role for Shf diffusion in Hh gradient formation. This is consistent with the observation that a membrane-bound form of Shf (Shf-CD8) is sufficient to rescue Hh signalling in *shf^EY^* null mutants when expressed in either the producing or receiving cells, or when expressed in cell populations away from the Hh receiving cells (Figure 7 and Supplementary Figure 10), and strongly supports cytoneme-mediated Hh transport.

### Role of Shf during Hh gradient formation

Both Ihog and Shf are needed to stabilise Hh at the cell membranes (Avanesov and Blair, 2013; Bilioni et al., 2013; Glise et al., 2005; Gorfinkiel et al., 2005; Yan et al., 2010; Yao et al., 2006). An interaction between Shf and Ihog has already been described (Avanesov and Blair, 2013; Bilioni et al., 2013), suggesting that both proteins collaborate in the process of extracellular Hh stabilization. In addition, AalphaFold modeling of Ihog and Shf simultaneously revealed a putative interaction, formed by the EGF domains of Shf and the Ig1/Ig4 domains of Ihog. This interaction is also supported by the behaviour of the shf2 allele with a point mutation in the EGF repeats (C-374-S) (Gorfinkiel et al., 2005; Glise 2005) (Figure 4). As predicted, we demonstrate both in vivo and in vitro that the Ig domains of Ihog interact with Shf (Figure 4). This interaction has been shown to be required for Hh stabilisation, as expressions of Ihog mutants, which individually cannot interact with Hh, are able to restore Hh accumulation in the P compartment when expressed in pairwise combination. A possible explanation for this complementation is that Hh would be able to interact with the Fn1 domain of Ihog as was previously described (McLellan et al., 2006; Simon et al., 2021; Yang et al., 2021; Zheng et al., 2010), but this interaction would not be sufficient for Hh to be correctly retained at membranes: for this retention the collaboration of Shf (which in turn interacts with the Ig domain) would be necessary.

We also observed that Ihog Ig domains play distinct roles on Hh stabilization and signal transduction. This is supported by the fact that the Ihog mutant protein pairwise combinations able to accumulate Hh were not able to transduce Hh signaling when expressed in the A compartment. These different behaviors of Ihog in each compartment could largely depend on the different affinity of Ihog-Ihog, Ihog-Hh and Ptc-Ihog-Hh interactions, as it has been proposed (Wu et al., 2019; Yang et al., 2021). In those lines, we have performed structural predictions that indicate that Shf interaction with Hh is lost by its interaction with the coreceptor complex formed by Ptc and Ihog (Supplementary Figure 8, asterisk). The most parsimonious model for these results is that a single Ihog molecule would transfer Hh from its complex with Shf and present it to Ptc, using the Ig and FN domains, respectively.

In addition to its role in stabilising Hh at the cell surface through its interaction with Ihog, Shf also facilitates the release of Hh from P compartment cells to form a gradient in the A compartment (Glise et al., 2005; Gorfinkiel et al., 2005). The Shf rescue of the shortening of the Hh gradient caused by Ihog overexpression (Supplementary Figure 12) indicates the effect of Shf on the Hh release process. In this context, we have also observed overexpressing Shf reverts the Hh accumulation in Dlp mutant cells (Figure 8). Masking the Hh lipids could enable Shf to act as a solubiliser for Hh exchange at the contact sites between the A- and P-compartment cytonemes. The involvement of Shf in the exchange of Hh between presenting and receiving cytonemes is also suggested by the observation that endogenous Shf and Hh are retained extracellularly and together in the cytonemes of Hh receiving cells following expression of morphotrap in the A compartment of GFP-tagged Shf flies. In support of a role for Shf in Hh solubilisation, structural modeling predicts that Shf is able to interact with Hh through both its EGF repeats and the WIF domain, in an interaction that is predicted to be dependent on the absence of Ihog in the complex (Supplementary Figure 11). In agreement with this hypothesis, the *shf^919^*mutant allele has a point mutation (C-638-S) (Supplementary Figure 11) located within the domain of interaction with Hh. This genetic evidence supports the existence of a functionally relevant Shf–Hh interaction in vivo.

### Model of Hh release and reception differences between Drosophila and vertebrates

We propose the following model for Hh release and reception: Hh reaches the basal side of the wing disc epithelium after undergoing intracellular recycling from the apical side, which involves Disp (Figure 9 step 1) (Callejo et al., 2011; Gradilla et al., 2014; Hall et al., 2021; Stewart et al., 2018). An aberrant trafficking of Hh in *disp* mutant cells could block the exit of Hh, which is in line with previous findings (Callejo et al., 2011; Gradilla et al., 2014; Hall et al., 2021; Stewart et al., 2018). Alternatively, Disp could be involved in the flipping of Hh to the outer membrane bilayer as it has been also proposed (Wang et al., 2021). In any case, both possibilities are in agreement with the observed intracellular localization of Hh in *disp* mutant cells (Figure 8). Once Hh is exposed extracellularly, it is stabilised by Ihog. As Ihog and Disp also interact (Callejo et al., 2011), Disp may deliver the morphogen to Ihog. Efficient Hh stabilization requires cooperation between Ihog and Shf, consistent with their direct interaction through the Ihog Ig domains. (Figure 9, step 2). In addition, the glypicans also collaborate through their HS chains in maintaining Ihog protein levels in the plasma membrane (Simon et al., 2021). Finally, the glypican Dlp and Shf intervene to facilitate the exit of lipidated Hh from the producer cells to reach the receptor cells (Callejo et al., 2011; Bilioni et al., 2013; Avanesov and Blair, 2013).

Morphogen exchange can occur at contact sites between Hh-producing and Hh-receiving cytonemes (Chen et al., 2017b; González-Méndez et al., 2017).This process involves Shf, which enables the mobilisation of Hh from the P cells, and may collaborate with Ihog in the uptake of Hh at the cytoneme contacts. (Figure 9, step 3). Therefore, it is not unreasonable to suppose that Ihog and Shf in the recipient cells, together with the competition of Ptc for Hh, releases Ihog-Shf-Hh-Ihog interactions for Hh reception. In agreement with this hypothesis, Shf has been shown to rescue *shf^EY^* null mutants when it is expressed attached to membranes (Shf-CD8) in the receiving cells. In addition, a different behavior of Ihog in each compartment could largely depend on the different affinity of Ihog-Ihog, Ihog-Hh and Ptc-Ihog-Hh interactions, as it has been proposed (Wu et al., 2019; Yang et al., 2021). Finally, the Hh receptor-complex, that in addition of Ihog, also involves the glypican Dlp (Desbordes and Sanson, 2003; Lum et al., 2003; Williams et al., 2010), could present the morphogen to its canonical receptor Ptc (Figure 9, step 4), initiating Hh signalling by SMO activation (Alcedo et al., 1996). Future work using improved endogenous tagging strategies and higher-sensitivity live imaging will be required to directly visualize the proposed handover of Hh–Shf complexes at cytoneme contact sites. Such approaches should help resolve the dynamics and molecular sequence of events underlying ligand exchange in vivo.

Cytoneme-mediated Hh signalling has also been described in mice (Hall et al., 2024) and in chicks (Sanders et al., 2013). Moreover, a similar model for SHH release and transfer has been proposed based on biochemical data in mammalian tissue culture cells (Tukachinsky et al., 2012; Wierbowski et al., 2020). This model implies the diffusible protein SCUBE2 that binds SHH, “hiding” its lipid modifications and facilitating the uptake of SHH by CDON/BOC. Then, SHH is transferred to GAS1, which in turn, presents the morphogen to its receptor PTCH1 (Wierbowski et al., 2020). Neither the SCUBE2 nor the GAS1 proteins are present in *Drosophila*. However, it is interesting to note that the SCUBE2 protein could mask SHH lipids in a manner similar to Shf, since both proteins are required for the extracellular stability of lipidated Hh only (Glise et al., 2005; Gorfinkiel et al., 2005). . Besides, considering that GAS1 also masks the lipid modification of SHH, as it has recently been described for Dlp in lipidated Wingless (McGough et al., 2020), the function of GAS1 as a SHH co-receptor is reminiscent of that described for Dlp in *Drosophila* (Desbordes and Sanson, 2003; Lum et al., 2003; Williams et al., 2010)

Shf is the ortholog of the human Wnt Inhibitory factor 1 (WIF1 protein) (Glise et al., 2005; Gorfinkiel et al., 2005; Hsieh et al., 1999). Unlike Shf, WIF1 regulates Wnt signalling (Hsieh et al., 1999). As Hh, Wnt ligands are also lipidated and it is also through the lipid moieties that WIF1 interacts with Wnt (Malinauskas, 2008; Malinauskas et al., 2011). Although originally identified as a Wnt inhibitor, WIF1 has more recently been proposed to function as a carrier, facilitating Wnt transport in the extracellular space (de Almeida Magalhaes et al., 2024). Given the growing evidence for cytoneme-mediated Wnt trafficking (Routledge and Scholpp, 2019; Stanganello et al., 2015; Zhang et al., 2024), it is plausible that WIF1 fulfills a role analogous to that of Shf in contact-dependent ligand delivery.

To date, diffusible modulators of morphogen distribution have been identified in most pathways, including Hh (Chuang and McMahon, 1999; Glise et al., 2005; Gorfinkiel et al., 2005; Tukachinsky et al., 2012), Wnt (de Almeida Magalhaes et al., 2024; Mihara et al., 2016; Mulligan et al., 2012), BMP (Ashe and Levine, 1999; Holley et al., 1996; Shimmi et al., 2005), and Nodal (Müller et al., 2012). These extracellular carriers have sometimes been seen as alternatives to direct, contact-mediated transport mechanisms such as cytonemes (Bilioni et al., 2013; Schlissel et al., 2024). Our findings provide the first evidence that morphogen carriers may also function during cytoneme-mediated transport, opening the door to a broader re-evaluation of how diffusible carriers operate within contact-dependent signaling systems. Together, our results support a model in which Shf functions not as a freely diffusing carrier but as a membrane-constrained cofactor that operates within the cytoneme-based Hedgehog transport pathway.

## Materials and methods

### *Drosophila* strains and husbandry

The description of mutations, insertions and transgenes is available at FlyBase (http://flybase.org). The following mutants and transgenic strains were used: *shf^EY^*(BDSC # 15899), *dlp^20^* (Franch-Marro et al., 2005), *disp^SH23^* (Amanai and Jiang, 2001), and *shf^2^* (BDSC # 112), *shf^919^*(Gorfinkiel et al 2005). *shf^GFP^* is an insertion of GFP in the reading frame of the endogenous *shf* locus (Aguilar et al., 2024b). Flies were kept at 25 °C unless specified.

The Gal4 and Gal80 lines used were: *hh*.Gal4 (Tanimoto et al., 2000), *ptc*.Gal4 (Hinz et al., 1994), *ap*.Gal4 (Calleja et al., 1996), *en*.Gal4 (with expression induction only in the posterior compartment, provided by Christian Dahmann), *ubx*.Gal4 (L. F. de Navas et al., 2006), AB1.Gal4 (BDSC #1824), *tub*.Gal4 (Lee and Luo, 1999), and tub.Gal80^ts^ (BDSC #7017, #7108).

The *UAS* lines used were: *UAS*.CD8-GFP (Lee and Luo, 1999), *UAS*.morphotrap (Harmansa et al., 2015), *UAS*.shf-V5 (Glise et al., 2005), *UAS*.ihog-RFP, *UAS*.ihogCT-RFP, *UAS*.ihogΔIg-RFP, *UAS*.ihogΔFN-RFP, *UAS*.ihogΔFn1-RFP and *UAS*.ihogΔFn1***:RFP (Simon et al., 2021), *UAS*.Disp:GFP (Callejo et al., 2011), *UAS*.*ihog* RNAi (VDRC #102602), *UAS*.*disp* RNAi (BDSC #44633), *UAS*.*dlp* RNAi (VDRC #10299), *UAS*.*dally*-RNAi (VDRC#14136), UAS-secGFP (this study), UAS-GFP-CD8 (this study) and UAS-Shf-CD8 (this study).

### Clonal analysis

Mutant clones were generated by heat shock-induced Flp-mediated mitotic recombination. Briefly, individuals grown at 17°C were incubated at 37 °C for 45 minutes 48–72 hours after egg laying (AEL). The genotypes in each case were: *y,w,hsFLP; UAS.CD8-GFP / tub.Gal4; FRT82B,tub.Gal80 / FRT82B,disp^SH23^ y,w,hsFLP; dlp^20^,FRT2A / tub.Gal80,FRT2A; tub.Gal4 / UAS.GFP y,w,hsFLP; UAS.CD8-GFP/UAS.shf:V5,tub.Gal4 ; RT82B,tub.Gal80 / FRT82B,disp^SH23^ y,w,hsFLP; FRT2A dlp^20^ tub.Gal80 / FRT2A ; UAS.shf:V5 / tub.Gal4, UAS.GFP*

### Cloning and transgenesis

*UAS*.Shf-CD8: The *shf* cDNA (DGRC stock 2344) was amplified with the primers 5’ AAAAAACTCGAGGTACCCGCCAGCAGCACCACAAT 3’ and 5’ AAAAAAGTATCGCATGCGAACTTGGAGTCATCGTTTC 3’ to introduce KpnI and SphI restriction sites. The product was subsequently subcloned into the pLOT-Morphotrap plasmid (Harmansa et al., 2015) in place of the antiGFP nanobody, in frame with the CD8 and mCherry sequences.

*UAS*.GFP-CD8: The GFP coding sequence (cds) was subcloned into the pLOT-morphotrap using the KpnI SphI enzyme pair. Subsequently, the sequence encompassing the signal peptide GFP and CD8 was amplified using the primers: 3’-AAAGCGGCCGCTTAGCTGTGGTAGCAGATGAGAGTGATGA-5’ and 3’-TAAAGATCTGCCACCATGGCCTCAC-5’. The fragment was subcloned into the pUASTattb plasmid using BglII and NotI restriction enzymes.

*UAS*.secGFP: The GFP cds, preceded by the CD8 signal peptide was amplified from a pLOT-GFP-CD8-mCherry plasmid using the primers: 3’-TTATTGAATACAAGAAGAGAACTCTGAATAGGGAATTGGGGTGCCACCATGGCCTCAC-5’ and 3’-atgtcacaccaCAGAAGTAAGGTTCCTTCACAAAGATCCTttacttgtacagctcgtccatgcc-5’. The pUASTattb was linearized using EcoRI and XbaI. The PCR product and the linearized plasmid were then assembled using Gibson assembly.

The transgenic Drosophila lines were generated by microinjection as previously described (Aguilar et al., 2024a). The acceptor ‘*landing-pads’* employed were M{3xP3-RFP.attP}ZH-22A (insertion into 2L-22A) and M{3xP3-RFP.attP}ZH-86Fb (insertion into 3R-86F). Injected lines also contained the transgene M{vas-int.Dm}zh-2A.

### Immunostaining of imaginal wing discs

Immunostaining was performed following standard protocols (Capdevila and Guerrero, 1994). Briefly, imaginal discs from third instar larvae were fixed in 4% paraformaldehyde in PBS for 20 minutes at room temperature (RT), followed by permeabilization in PBT (PBS with 0.1% Triton X-100) for 30 minutes at RT. Samples were then blocked for 1 hour at RT in PBT supplemented with 1% BSA, and subsequently incubated overnight at 4 °C with the primary antibody diluted in the same buffer. The following day, samples were washed in PBT for 15 minutes at RT, then incubated for 1 hour at RT with the appropriate fluorescent secondary antibodies (ThermoFisher) diluted 1:400 in PBT. This was followed by three 15-minute washes in PBT and two final 15-minute washes in PBS at RT. Samples were mounted on a glass slide in Vectashield mounting media, covered with a mounting glass and sealed with colorless nail-polish. The primary antibodies used and the concentration at which they were used are indicated in the corresponding table in the Materials and Methods section.

For extracellular immunohistochemistry, samples were dissected in M3 Drosophila tissue culture medium and directly incubated in the primary antibodies diluted in M3 for 1h at RT. Samples were then thoroughly washed in the same buffer prior to fixation with 4% PFA in M3. The primary antibodies were used at concentrations four times higher than those indicated for normal staining in the table in the Materials and Methods section. After fixation, samples were permeabilized and followed the same protocol as the standard immunostaining.

### Microscopy and image processing of wing imaginal discs

The following equipment was used for the acquisition of immunofluorescence images: LSM900 Laser Scanning Confocal Microscope coupled to an AxioImager 2 upright microscope (Zeiss). LSM880 Laser Scanning Confocal Microscope with an AiryScan unit. LSM710 Laser Scanning Confocal Microscope coupled to an AxioImager.M2 upright microscope (Zeiss). LSM800 Laser Scanning Confocal Microscope coupled to an Axio Observer inverted microscope (Zeiss). For each experiment, at least five images of imaginal discs were acquired in three independent experiments. The minimum resolution of each image was at least 1024x1024 pixels. Images of Drosophila wings were acquired using an AxioImager M1 upright microscope (Zeiss) coupled to a DMC6200 camera (Leica).

### Quantification method and numerical analysis of Hh and Shf recruitment

Fluorescence quantification was performed in FIJI/ImageJ. For each imaginal disc, regions of interest (ROIs) of identical size were placed over the experimental region and over an internal control region within the same disc. Mean fluorescence intensity was measured for each ROI, and the experimental value was normalized to the control ROI from the same disc to obtain a normalized fluorescence ratio for each sample. Statistical analyses were performed in GraphPad Prism. Differences among multiple groups were assessed using one-way ANOVA followed by Tukey’s multiple-comparisons test. The significance threshold was set at α = 0.05.

### Quantification of the apicobasal profiles of intensity and of the gradients of Hh signaling

The apicobasal profiles of immobilized Hh:GFP and Shf:GFP were obtained as follows. First, high resolution Z-stacks were acquired using a point scanning confocal. In imageJ, each image was cropped to a ROI of consistent size placed in the middle of the wing pouch, covering the central stripe. A sagittal Z-stack was obtained using the re-slice tool. This sagittal Z stack was then projected using the sum of intensities of all planes, channels were split into different images. To limit the downstream quantification to only the immobilized ligands, the morphotrap channel was used to generate a mask of the receiving cells using the threshold tool. The mask was then used to clip the GFP channel using the image calculator tool, combining both images via the AND function. The intensity profile was then extracted using the “Plot profile” command. Mean values and standard deviations of several discs were then plotted using GraphPad’s Prism software.

### Structure modelling

To perform in silico protein structure prediction, we used AlphaFold 3 (Abramson et al., 2024) via the AlphaFold server. Each prediction was run with the default number of recycles to generate five relaxed models, which were ranked according to their predicted template modeling (pTM) scores. All five models were analyzed, and the highest-confidence prediction was selected for representation. Predicted alignment error (PAE) plots were generated and visualized in ChimeraX. Unless otherwise stated, protein models included N-acetyl glycosylation at residues annotated in the UniProt database. For Hh, only the N-terminal peptide was modeled, including palmitoylation but omitting the Cholesterol moiety. Structural representations were prepared in ChimeraX. Pseudobonds in Figure 4C were generated using the “alphafold contacts” command in ChimeraX to visualize inter-residue PAE values, colored by error magnitude and filtered by spatial distance and confidence thresholds.

### Co-Immunoprecipitation and Western-blot analysis

Protein extraction was performed from extracts of third-instar Drosophila melanogaster salivary glands. To achieve this, transgenes under the control of the *UAS* sequence were expressed using the salivary gland AB1.Gal4 driver. For salivary gland tissue extraction, third-instar larvae were collected and dissected, and the salivary glands were stored at -80 °C. The number of salivary glands used in each protein extraction was 50 pairs of glands.

Protein immunoprecipitation was carried out from the protein extracts using a magnetic matrix (Dyneabeads) to which an antibody against RFP (ChromoTek RFP-Trap® Magnetic Agarose) was bound. The elution of immunoprecipitated proteins was performed by removing the RIPA buffer and adding 25 μl of loading buffer (Laemmli buffer), heating the mixture for 5 minutes at 95 °C, and collecting the supernatant after placing the Eppendorf tube in the DynaMag™-2 magnet. The samples were re-suspended in a sample-buffer with β-mercaptoethanol and subjected to 1D SDS-Page (8%) and Western blotting. Blotted membranes were probed with the primary antibodies and with secondary antibodies for each case (see table of Material and Methods), and imaged with the 364-Odyssey equipment.

---

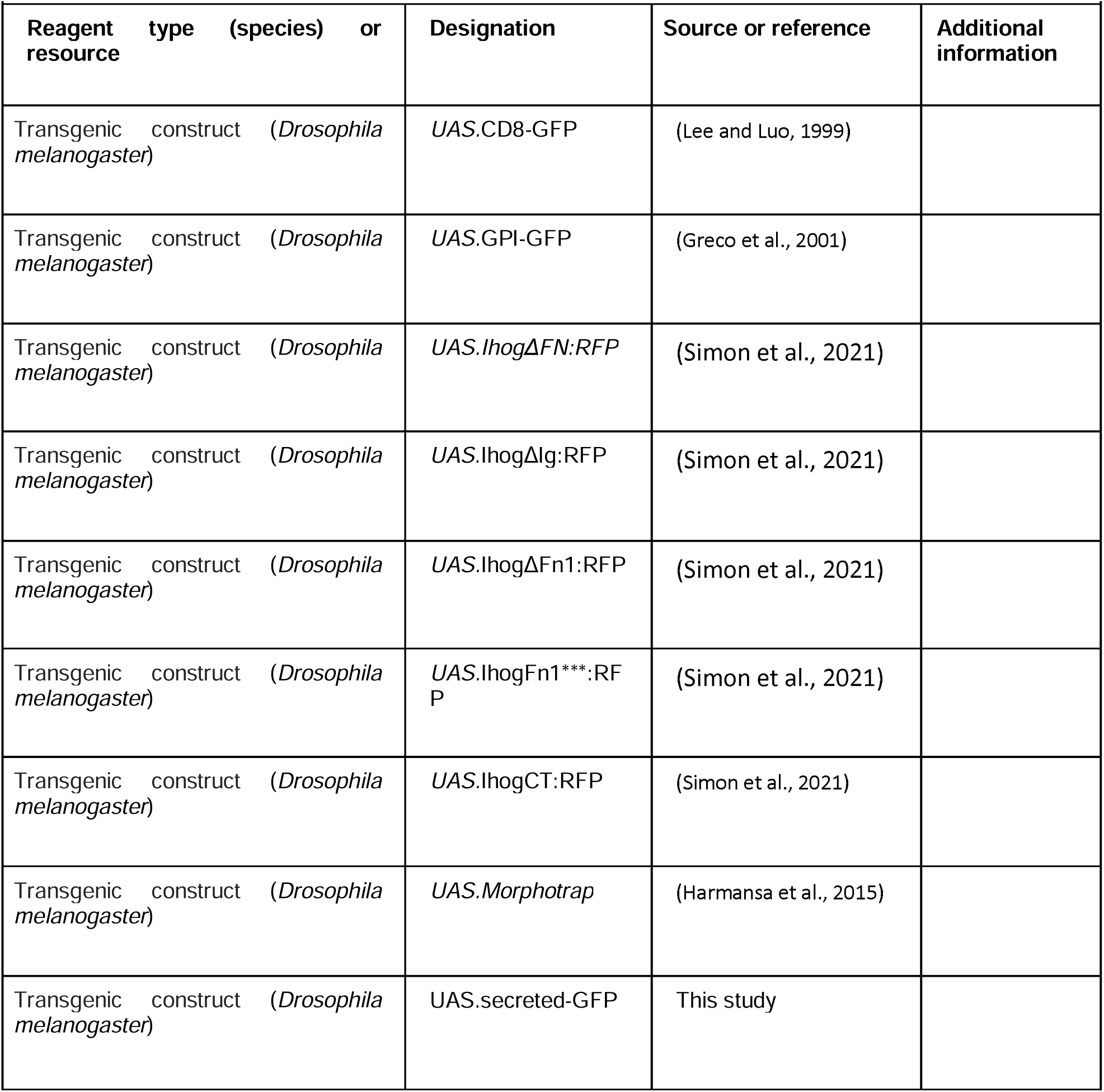
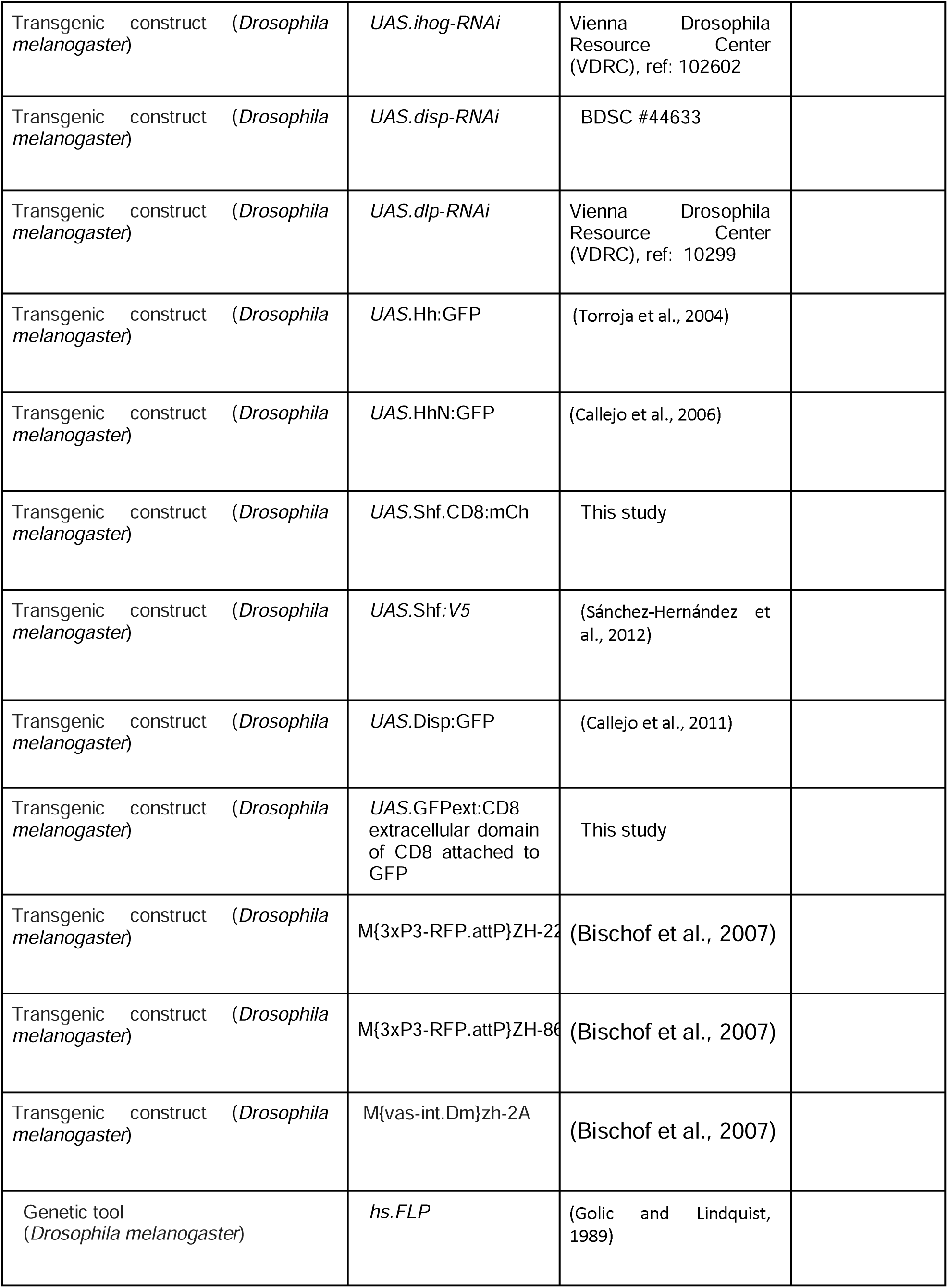
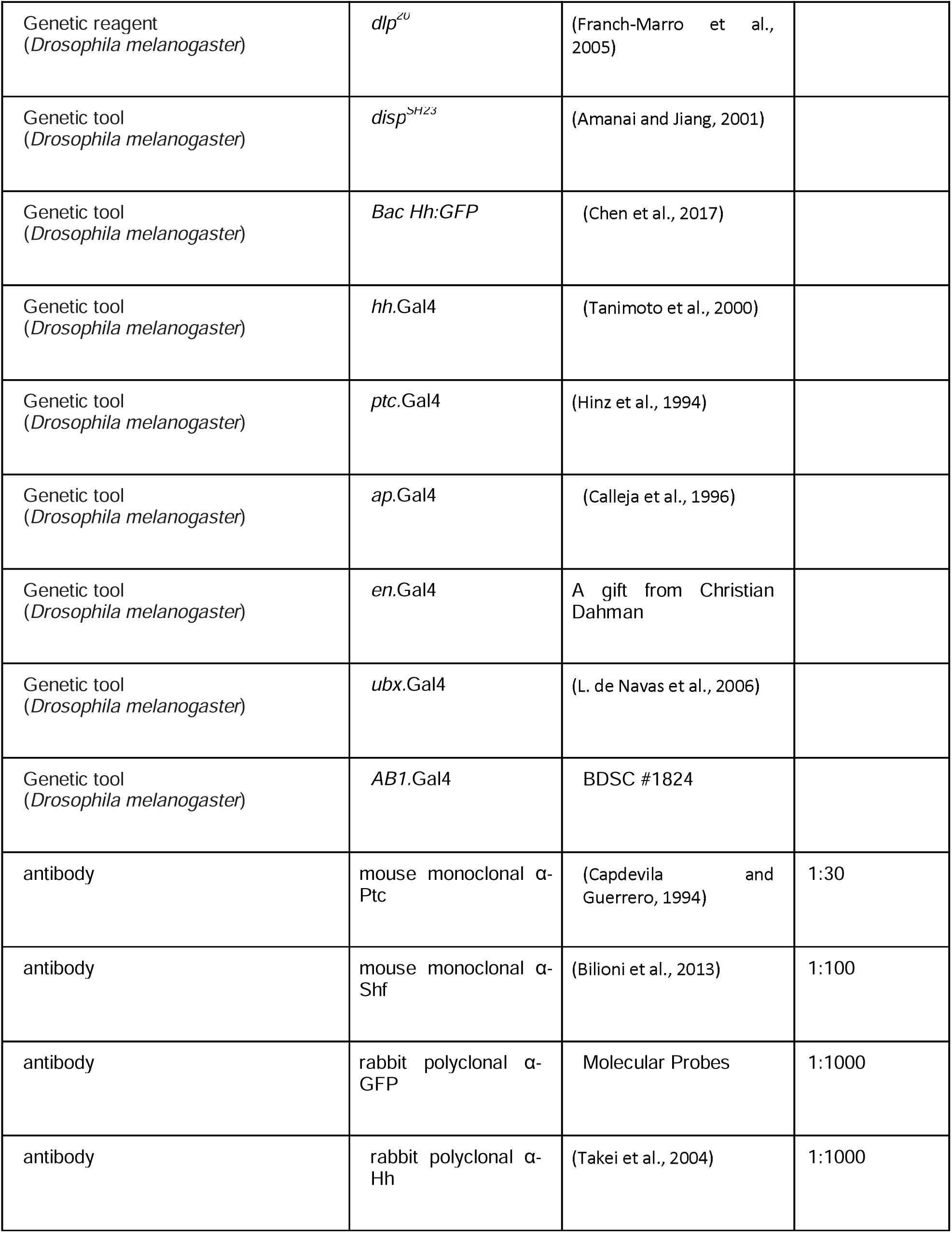
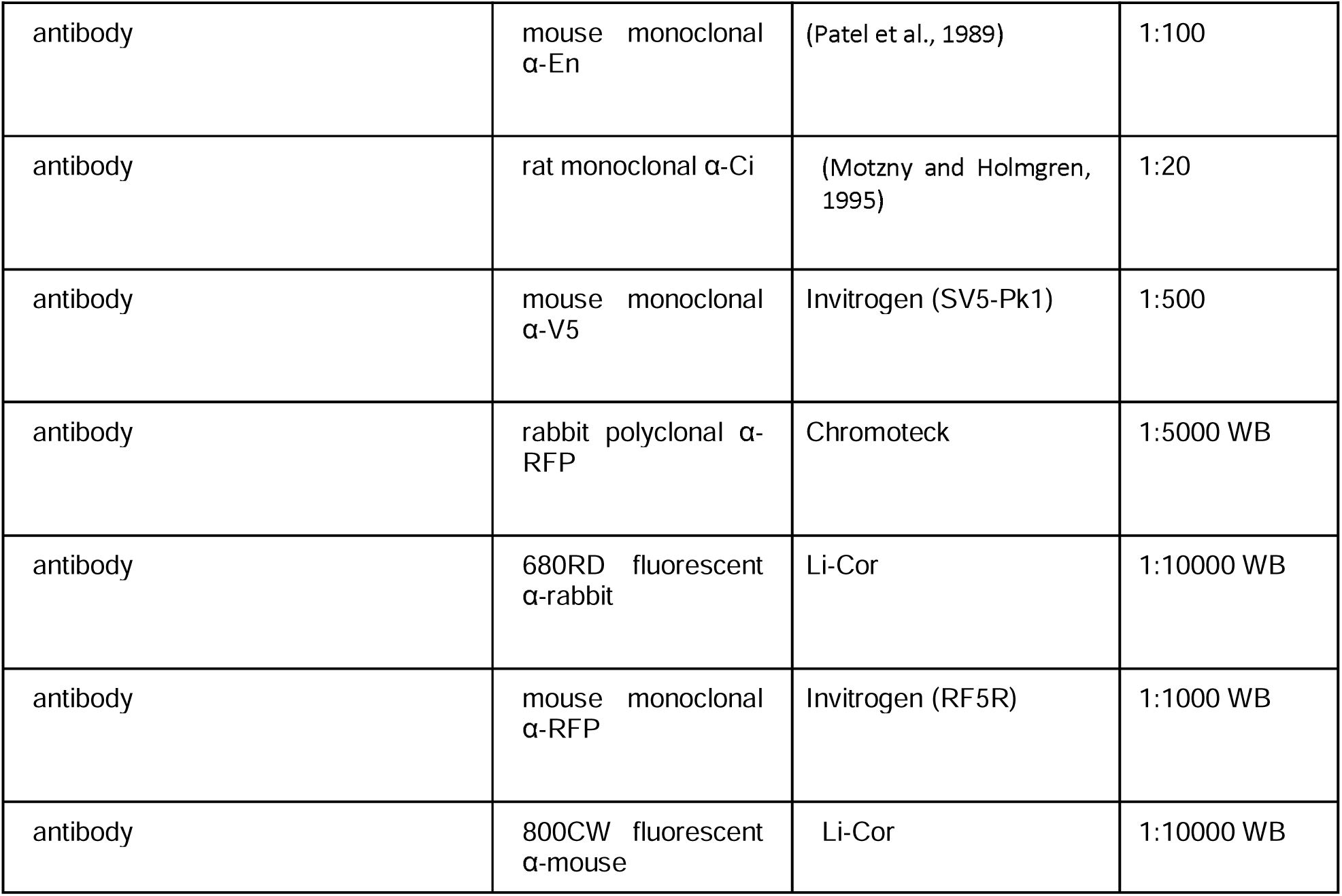
Key Resources Table.

| Key Resources Table |  |  |  |  |
| --- | --- | --- | --- | --- |
| Reagent type (species) or resource | Designation | Source or reference | Additional information |  |
| Transgenic construct (*Drosophila melanogaster*) | UAS.CD8-GFP | (Lee and Luo, 1999) |  |  |
| Transgenic construct (*Drosophila melanogaster*) | UAS.GPI-GFP | (Greco et al., 2001) |  |  |
| Transgenic construct (*Drosophila melanogaster*) | UAS.*lhog*ΔFN:RFP | (Simon et al., 2021) | | |
| Transgenic construct (*Drosophila melanogaster*) | UAS.*lhog*Δlg:RFP | (Simon et al., 2021) | | |
| Transgenic construct (*Drosophila melanogaster*) | UAS.*lhog*ΔFn1:RFP | (Simon et al., 2021) | | |
| Transgenic construct (*Drosophila melanogaster*) | UAS.*lhog*Fn1<sup>***</sup>:RFP | (Simon et al., 2021) |  |  |
| Transgenic construct (*Drosophila melanogaster*) | UAS.*lhog*CT:RFP | (Simon et al., 2021) |  |  |
| Transgenic construct (*Drosophila melanogaster*) | UAS.*Morphotrap* | (Harmansa et al., 2015) |  |  |
| Transgenic construct (*Drosophila melanogaster*) | UAS.secreted-GFP | This study |  |  |

|  |  |  |
| --- | --- | --- |
| Transgenic construct (*Drosophila melanogaster*) | *UAS.ihog-RNAi* | Vienna Drosophila Resource Center (VDRC), ref: 102602 |
| Transgenic construct (*Drosophila melanogaster*) | *UAS.disp-RNAi* | BDSC #44633 |
| Transgenic construct (*Drosophila melanogaster*) | *UAS.dlp-RNAi* | Vienna Drosophila Resource Center (VDRC), ref: 10299 |
| Transgenic construct (*Drosophila melanogaster*) | *UAS.Hh:GFP* | (Torroja et al., 2004) |
| Transgenic construct (*Drosophila melanogaster*) | *UAS.HhN:GFP* | (Callejo et al., 2006) |
| Transgenic construct (*Drosophila melanogaster*) | *UAS.Shf.CD8:mCh* | This study |
| Transgenic construct (*Drosophila melanogaster*) | *UAS.Shf:V5* | (Sánchez-Hernández et al., 2012) |
| Transgenic construct (*Drosophila melanogaster*) | *UAS.Disp:GFP* | (Callejo et al., 2011) |
| Transgenic construct (*Drosophila melanogaster*) | *UAS.GFPext:CD8*<br>extracellular domain of CD8 attached to GFP | This study |
| Transgenic construct (*Drosophila melanogaster*) | M{3xP3-RFP.attP}ZH-22 | (Bischof et al., 2007) |
| Transgenic construct (*Drosophila melanogaster*) | M{3xP3-RFP.attP}ZH-86 | (Bischof et al., 2007) |
| Transgenic construct (*Drosophila melanogaster*) | M{vas-int.Dm}zh-2A | (Bischof et al., 2007) |
| Genetic tool (*Drosophila melanogaster*) | *hs.FLP* | (Golic and Lindquist, 1989) |

|  |  |  |  |
| --- | --- | --- | --- |
| Genetic reagent<br>(*Drosophila melanogaster*) | *dlp*<sup>20</sup> | (Franch-Marro et al., 2005) |  |
| Genetic tool<br>(*Drosophila melanogaster*) | *disp*<sup>SH23</sup> | (Amanai and Jiang, 2001) |  |
| Genetic tool<br>(*Drosophila melanogaster*) | *Bac Hh:GFP* | (Chen et al., 2017) |  |
| Genetic tool<br>(*Drosophila melanogaster*) | *hh.Gal4* | (Tanimoto et al., 2000) |  |
| Genetic tool<br>(*Drosophila melanogaster*) | *ptc.Gal4* | (Hinz et al., 1994) |  |
| Genetic tool<br>(*Drosophila melanogaster*) | *ap.Gal4* | (Calleja et al., 1996) |  |
| Genetic tool<br>(*Drosophila melanogaster*) | *en.Gal4* | A gift from Christian Dahman |  |
| Genetic tool<br>(*Drosophila melanogaster*) | *ubx.Gal4* | (L. de Navas et al., 2006) |  |
| Genetic tool<br>(*Drosophila melanogaster*) | *AB1.Gal4* | BDSC #1824 |  |
| antibody | mouse monoclonal α-Ptc | (Capdevila and Guerrero, 1994) | 1:30 |
| antibody | mouse monoclonal α-Shf | (Bilioni et al., 2013) | 1:100 |
| antibody | rabbit polyclonal α-GFP | Molecular Probes | 1:1000 |
| antibody | rabbit polyclonal α-Hh | (Takei et al., 2004) | 1:1000 |

|  |  |  |  |
| --- | --- | --- | --- |
| antibody | mouse monoclonal α-En | (Patel et al., 1989) | 1:100 |
| antibody | rat monoclonal α-Ci | (Motzny and Holmgren, 1995) | 1:20 |
| antibody | mouse monoclonal α-V5 | Invitrogen (SV5-Pk1) | 1:500 |
| antibody | rabbit polyclonal α-RFP | Chromotek | 1:5000 WB |
| antibody | 680RD fluorescent α-rabbit | Li-Cor | 1:10000 WB |
| antibody | mouse monoclonal α-RFP | Invitrogen (RF5R) | 1:1000 WB |
| antibody | 800CW fluorescent α-mouse | Li-Cor | 1:10000 WB |

## Acknowledgements

We are grateful to the late Dr. Pedro Ripoll, whose daily interactions had a critical impact on the development of this story and that of the authors of this manuscript. We would also like to thank Dr. Nicole Gorfinkiel for comments on the manuscript, to Eleanor Simon for her help with the live imaging of Supplementary movie 1 and to Lara Rodenstein for her excellent technical assistance. We are also grateful to the imaging core facilities of the Biozentrum and the Centro de Biología Molecular for their technical assistance and equipment contribution. Finally, we are also thankful to P. Beachy, J. Treisman, R. Holgreen, T. Tabata, S. Eaton and T. Kornberg for stocks and reagents. The work at IG laboratory was supported Grant PID2020-114533GB-C21 to IG from the Spanish Ministry of Science, Innovation and Universities with fellowship PRI 2018-085510 for CJJ and a fellowship PRI 2021-097741 for CFP, and by institutional grants from the Fundación Areces and from Banco de Santander to the CBMSO. The work in the laboratory of M.A. was supported by grants from the Swiss National Science Foundation (310030_192659/1) and by funds from Kanton Basel-Stadt and Basel-Land. GA was supported by ‘Fellowships for Excellence’ from the International PhD Program in Molecular Life Sciences of the Biozentrum, University of Basel.

## Supplementary figures

**Supplementary Figure 1. High-magnification views of basal contacts revealed by membrane-tethered ligands.**

A–A’’) Wing disc expressing Morphotrap in Hh-receiving cells (dpp.LexA) and GFPext-CD8 in Hh-producing cells (hh.Gal4). A) Scheme of expression domains. A’) Basal view showing GFPext-CD8 accumulation at the A/P interface. A’’) Magnified view highlighting thin basal protrusions (arrows) emanating across the compartment boundary. B–B’’) Same experimental configuration using GPI:GFP. B) Scheme of expression domains. B’) Basal view showing GPI:GFP accumulation at the interface. B’’) Magnified view (arrows) revealing a network of basal contacts with morphology comparable to that observed with GFPext-CD8.

**Supplementary Figure 2. Summary schemes of retention patterns for synthetic ligands.**

Schematic overview of the immobilization behaviors observed in Figure 1 for ligands with different membrane tethering properties. No membrane tethering (Secreted GFP): the ligand disperses broadly and is captured uniformly along the apical, lateral, and basal membranes of morphotrap-expressing cells. Strong membrane tethering (GFPext-CD8): the ligand accumulates preferentially at cell–cell contact sites and is enriched within basal protrusive contacts spanning both compartments. Intermediate membrane tethering (GPI:GFP): basal protrusions are preferentially stabilized toward the P compartment, while a substantial fraction of signal remains associated with the lateral and apical membranes of the first receiving cell rows.

**Supplementary Figure 3. Hh:GFP immobilisation by the morphotrap.**

A) Close-up in basal sections of cytonemes loaded with Hh:GFP upon expression of Morphotrap (red) in the A compartment using the *ptc*.Gal4; *tub*.Gal80ts driver for 24h. A’) Enlargement of the cropped area in A to show the colocalisation of Hh:GFP and the Morphotrap in round structures in some places along cytoneme length (arrowheads). B) Close up of cytonemes protruding from a Morphotrap-expressing clone situated close to the Hh:GFP source. Notice the high levels of Hh:GFP accumulated in cytonemes and in the enlarged structures.

**Supplementary Figure 4. Model for Ihog and Boi interactions with Shf**

A) and A’) Structural prediction of the globular domains of Ihog and their intramolecular interactions. Notice the robust prediction of the closed conformation of the four Ig domains. B) and B’) Structural prediction of the Boi-Shf complex. Notice the conservation of the structural features between boy and Ihog (Panel A) as well as the conserved interaction surface of Boi-Shf and Ihog-Shf (Figure 5A) mediated by the EGF domains of Shf and the Ig domains of Ihog.

**Supplementary Figure 5. Hh recruitment by Ihog.**

A) Hh accumulation (visualized using anti-Hh antibody) in wing discs caused by overexpression of different Ihog mutant forms using the *ap.Gal4; tub.Gal80^ts^* driver; the ventral domain serving as an endogenous control: *UAS.Ihog*Δ*FN:RFP* ; *UAS.IhogFn1***:RFP; UAS.Ihog*Δ*Fn1:RFP*. As can be observed, none of these Ihog constructs, in which the Hh interacting domain (Fn1 domain) is affected, are able to recruit Hh when overexpressed (Simon et al., 2021). (D-F) Build-up of Hh protein caused by co-overexpressing:*UAS.Ihog*Δ*Ig:RFP/UAS.IhogFn1***:RFP, UAS.Ihog*Δ*Ig:RFP/AS.Ihog*Δ*Fn:RFP*, and *UAS.Ihog*Δ*Fn:RFP/ UAS.IhogFn1***:RFP* using the same *ap.Gal4; tub.Gal80ts* driver. Note that the expression of those Ihog mutant combinations that do not meet the "complementary" condition for interactions with Hh, do not accumulate Hh.

**Supplementary Figure 6.** E**c**topic **Ihog constructs repress endogenous Ihog levels**

Basal sections of wing discs carrying an Ihog-GFP BAC transgene (containing the endogenous regulatory and coding sequences) showing endogenous Ihog levels under different overexpression conditions in the apterous (ap) domain. Top panels display the merged red (Ihog:RFP constructs) and green (Ihog-GFP BAC) channels. Bottom panels show the Ihog-GFP BAC signal alone (grayscale). Overexpression of Ihog:RFP (WT), IhogΔFn1:RFP, IhogΔIg:RFP, and related Ihog variants results in a strong reduction of endogenous Ihog-GFP signal within the ap domain compared to surrounding tissue.

**Supplementary Figure 7. Effects on Shf accumulation in stabilized cytonemes and on Hh signaling after expressing the Ihog mutant forms in the Hh receiving cells.**

A, B) Shf:GFP accumulation in Ihog stabilised cytonemes in a wing disc after Ihog.RFP (A) or IhogFn1*** (B) expression in Hh-receiving cells using the ptc-Gal4 driver. Notice the accumulation of Shf:GFP in both cases, despite the fact that Hh is known not to attach to cytonemes when IhogFn1*** is expressed (Simon et al., 2021). C) *ap*.Gal4 LexA>*LexAop*.Gal80, *ptc*.Gal4; *tub*.Gal80ts wing discs co-expressing different *UAS*.Ihog mutant forms in the ventral compartment. Note that the expression of IhogDFn1 alone and the coexpression of IhogDIg + IhogFn1*** or IhogDIg:RFP + IhogDFn1 have dominant negative effects on Hh signalling, which is reported by the high threshold Hh pathway targets Ptc and En.

**Supplementary Figure 8. Model of the Ptc-Hh-Shf-Ihog complex.**

A) Structural prediction of the Ptc-Hh-Shf-Ihog complex. Only globular domains are shown. Approximated areas of interaction are indicated. B) AlphaFold confidence matrix indicating the predicted aligned error of the model in A. Full length proteins were used for the prediction. Different proteins are indicated in colors. Notice that the interaction between Hh and Shf is abrogated in this complex (asterix). The last values of the matrix are dedicated to the PTMs.

**Supplement Figure 9. Hh signalling in *shf^EY^* null mutant wing disc is rescued by wild type Shf but not by Shf-CD8 when they are expressed in the PM.**

A) Drawing of a wing disc showing in a transversal section the two disc epithelia: Peripodial membrane (in green) and the disc proper (in grey). A scheme representing the two forms of Shf (wild type Shf and Shf-CD8) expressed in the peripodial membrane. B-B’’) Hh signalling in a *shf^EY^* null mutant wing disc visualised by the expression of the reporters Ptc (B) and Ci (B’), and the adult *shf^EY^*null mutant wing phenotype (B’’). (C-D’’) Effects on Hh signalling when expressing the *UAS*.Shf:V5 (C) and *UAS*.Shf-CD8:mCh (D) forms in a *shf^EY^*mutant background under the control of the *ubx*.Gal4 driver, which directs expression to the peripodial membrane of the wing disc. Only wild-type Shf expression in the peripodial membrane (C-C’’) is able to rescue Hh signalling in the disc proper epithelium (C’) and in the adult wing (C’’) of shf^EY^ mutants. In contrast, expression of Shf:CD8 (D) fails to rescue both Hh signalling (D’) and the wing phenotype (D’’) of shf^EY^ mutants.

**Supplementary Figure 10. Rescue of Hh signalling in *shf^EY^*mutants upon expression of Shf-CD8:mCh using the *magu*.Gal4 line.** A-A’’) Visualisation of the *magu* expression domain and the expression of the Hh signalling reporters Ci and Ptc by the induction of the *UAS*.CD8:RFP using the *magu*.Gal4 driver in a wild-type disc. B-C’’) Effects on Hh signalling upon expression of the UAS.Shf-CD8:mCh using the *magu*.Gal4 driver in a *shf^EY^* mutant wing imaginal disc (B) and upon expression of the UAS.Shf-CD8:mCh (B-B’’) and by simultaneously expressing an RNAi against SCAR using the same driver (C-C’’). D) Expression of Ptc and Ci in a *shf^EY^*mutant wing disc. It can be seen that the expression the Shf-CD8:mCh construct largely rescues the high Hh signalling values. On the other hand, by reducing the number and extent of cytonemes by the expression of the SCAR RNAi, the rescue of Hh signalling is much lower (C‘, C’’). The enlarged areas (yellow dotted line boxes) are 20x50 μm in size for Ptc and 20x100 μm for Ci.

**Supplementary Figure 11. Shf can interact with Hh via its EGF domains and WIF domain simultaneously.**

A) and A’) Structural prediction of Shf and Hh. The approximated surface of interaction is indicated. Only globular domains are shown in A for illustration purposes. Full proteins were employed for the prediction (See panel A’). B) Detail of the interaction between Hh and the EGF domains of Shf. In Blue-Orange Palette, the confidence of the interaction is measured using ChimeraX.

**Supplementary Figure 12. Shf can normalise the Hh retention of Hh and Hh signaling mediated by Ihog overexpression.**

A) *hh*.Gal4 tub.Gal80ts>Ihog:CFP wing disc after 22 hours at the restricted temperature and stained for the Hh pathway reporters Ptc and Ci. B) *hh*.Gal4 tub.Gal80ts>Ihog:CFP *UAS.*Shf- CD8:mChr wing disc after 22 hours at the restricted temperature and stained for Ptc and Ci. C) *hh*.Gal4 tub.Gal80ts>Ihog:CFP / UAS.Shf:CD8 wing disc after 22 hours at the restricted temperature and stained for Ptc and Ci. D) Quantification of Ptc expression gradient in each experimental condition. E) Quantification of Ci expression gradient in each experimental condition. Note the reduction of Ptc and Ci by the expression of Ihog:CFP and the normalized Ptc and Ci after coexpressing *UAS*.Ihog:CFP either *UAS*.Shf-CD8:mCh (B) or *UAS*.ShfV5 (C). F-F’’) *ap*.Gal4 tub.Gal80ts>Ihog:RFP wing disc after 22 hours at the restricted temperature and stained for Hh. Note the accumulation of Hh in the Ihog:RFP expressing cells. G-G’’) *ap*.Gal4 tub.Gal80ts>Ihog:CFP / UAS.Shf:CD8 wing disc after 22 hours at the restricted temperature and stained for Hh. Notice the decrease in Hh accumulation when both Ihog:RFP and UAS.Shf:CD8 are coexpressed.

**Supplementary Figure 13. Hh accumulation in *dlp^-/-^*and in *disp^-/-^* clones and after over expressing Shf.**

(A-B) Hh accumulation (visualized using specific anti-Hh antibody) in *dlp^-/-^* (A) and *disp^-/-^* (B) mutant clones labeled by expressing a nuclear GFP (A) and *UAS*.CD8:GFP (B) respectively. Note that in *dlp^-/-^* mutant cells there is a non-autonomous effect that rescues Hh accumulation at the periphery of the mutant territory (A’’, white arrowheads) that in the case of *disp^-/-^*the Hh accumulation is autonomous.

**Supplementary Figure 14. Analysis of functional epistasis among key proteins in the Hh secretion process.**

(A-C) Effect on Hh levels after expression of *UAS.disp RNAi* (A), *UAS.ihog RNAi* (B) and *UAS.dlp RNAi* (C) using *ap.Gal4* driver. (D-E) Effect on Hh levels after simultaneous expression of *UAS.disp RNAi/UAS.ihog RNAi* (D) and *UAS.dlp RNAi/UAS.ihog RNAi* (E) using the *ap.Gal4* driver. Note that the effect on Hh upon Disp elimination prevails over that of Ihog (D), and the effect upon Ihog elimination prevails over that of Dlp (E).

## Supplementary movies

**Supplementary Movie 1. Morphotrap dynamic behaviour of cytoneme-like structures in the pupal abdomen.**

Time-lapse imaging of abdominal histoblasts in a fly pupa expressing Morphotrap under the control of *ptc*.Gal4. Images are shown as maximum projections of multiple z-planes. Frames were acquired every 2 minutes. The movie illustrates the highly dynamic behaviour of these structures, which extend from Morphotrap-expressing cells.

**Supplementary Movie 2. Morphotrap stabilizes Hh-positive extensions in an BAC.*hh*:GFP background.**

Time-lapse imaging of abdominal histoblasts expressing Morphotrap under *ptc*.Gal4 in an BAC.*hh*:GFP background. Images are shown as maximum projections of multiple z-planes, acquired every 2 minutes. In contrast to the dynamic behaviour observed in Supplementary Movie 1, the extensions form a largely static network with little or no movement. The protrusions display multiple thickenings and contain numerous puncta positive for both GFP and Morphotrap signals.

**Supplementary Movie 3. Morphotrap stabilizes Shf-positive extensions in a *shf*:GFP background.**

Time-lapse imaging of abdominal histoblasts expressing Morphotrap under *ptc*.Gal4 in a Shf:GFP (endogenous *shfGFP* allele) background. Images are shown as maximum projections of multiple z-planes, acquired every 2 minutes. Similar to the Hh:GFP condition, the extensions form a largely static network with little or no movement and display multiple thickenings containing puncta positive for both GFP and Morphotrap signals.

